# Multi-scale structural alterations of the thalamus and basal ganglia in focal epilepsy as demonstrated by 7T MRI

**DOI:** 10.1101/2022.11.01.514655

**Authors:** Roy AM Haast, Benoit Testud, Julia Makhalova, Hugo Dary, Alexandre Cabane, Arnaud Le Troter, Jean-Philippe Ranjeva, Fabrice Bartolomei, Maxime Guye

## Abstract

Focal epilepsy is characterized by repeated spontaneous seizures that originate from cortical epileptogenic zone networks (EZN). More recently, analysis of intracerebral recordings showed that subcortical structures, and in particular the thalamus, play an important role in facilitating and/or propagating epileptic activity. This supports previously reported structural alterations of these structures. Nonetheless, between-patient differences in EZN (e.g., temporal vs. non-temporal lobe epilepsy) as well as other clinical features (e.g., number of epileptogenic regions) might impact the magnitude as well as spatial distribution of subcortical structural changes. Here we used 7 Tesla MRI T_1_ data to provide a comprehensive description of subcortical morphological (volume, tissue deformation, and shape) and longitudinal relaxation (T_1_) changes in focal epilepsy patients to evaluate the impact of the EZN and patient-specific clinical features. Our results showed widespread morphometric and T_1_ changes. Focusing on the thalamus, atrophy varied across nuclei but appeared most prominent for the TLE group and the ipsilateral side, while shortening of T_1_ was observed for the lateral thalamus, in particular. Multivariate analyses across thalamic nuclei and basal ganglia showed that volume acted as the dominant discriminator between patients and controls, while (posterolateral) thalamic T_1_ measures looked promising to further differentiate patients based on EZN. In particular, the observed differences in T_1_ changes between thalamic nuclei indicated differential involvement of thalamic nuclei based on EZN. Finally, the number of epileptogenic regions was found to best explain the observed variability between patients. To conclude, this work revealed multi-scale subcortical alterations in focal epilepsy as well as their dependence on several clinical characteristics. Our results provide a basis for further, in-depth investigations using (quantitative) MRI and SEEG data and warrant further personalization of intervention strategies, such as deep brain stimulation, for treating focal epilepsy patients.

## Introduction

Focal epilepsy is a chronic neurological disorder that is characterized by repeated spontaneous seizures caused by abnormal neuronal rhythms. These seizures typically start in one or several specific regions of the brain (i.e., epileptogenic zone, or EZ) and propagate to other regions or the whole brain. In approximately 30% of patients, epileptic seizures cannot be controlled by medication^1,2^, thereby significantly hampering the patient’s quality of life^3^. While a traditional view was to consider the EZ as a focal area of the cortex^4^, it is now recognized that focal epilepsy is a network disorder characterized by a hierarchical organization and anatomical specificity of subnetworks involved in the generation (i.e., epileptic zone network, EZN) and propagation (i.e., propagation network, PN) of epileptic activities across the brain^5^. Brain abnormalities might extend well beyond the EZ, involving regions that correlate with cognitive impairment but are only affected by seizure propagation or even spared from ictal discharges (i.e., non-involved networks).

Network approaches – based on the mapping of inter-regional connectivity or covariance – showed that focal epilepsy indeed affects the whole-brain and that focal structural and functional changes within the EZ appear to act more as a set of nodes in a larger network of affected regions^6,7^. For example, changes in cortical morphometry and connectivity have been consistently observed across the cortical mantle^8–12^. Direct longitudinal relaxation time (T_1_) mapping showed a prolongation of T_1_ relaxation times in temporopolar, orbitofrontal and para-hippocampal regions indicating alterations at the microstructural level as well^13^. Nonetheless, most of the published work focused on a single type of focal epilepsy and relatively little work has been done on comparing between patients characterized by different anatomically-defined EZ networks (EZN).

Moreover, increasing evidence points to a crucial role for subcortical structures in facilitating and/or propagating epileptic activity^14^. They play an important role in brain functioning as integrative hubs for brain connectivity^15^. In particular the thalamus, with its complex organization and nuclei, is a key gateway for cortico-subcortical interactions facilitating precise motor-sensory to broad cognitive functioning^16,17^. Therefore and not surprisingly, accumulating evidence using stereoelectroencephalography (SEEG)^14,18,19^ and MRI^18,20,21^ reveal a prominent involvement of the thalamus in TLE. Nonetheless, the amount of evidence compared to that focused on the cortex is relatively sparse and a comprehensive quantitative description of the thalamus and basal ganglia characteristics is lacking. For example, it remains elusive whether patterns of morphological and microstructural abnormalities across distinct, subcortical regions are (i) specific for the topographic organization of the EZN and/or can be linked to specific clinical variables including seizure frequency, age at onset and duration of epilepsy and presence of MRI-visible epileptogenic lesions.

To answer these questions, we will make use of ultra-high field (UHF, ≥7 Tesla) strength anatomical MRI data acquired in both temporal lobe epilepsy (TLE) and non-temporal lobe epilepsy (NTLE) patients. Growing amounts of evidence prove the clinical benefits of MRI at UHF strengths^22,23^. Compared to conventional 1.5 Tesla (1.5T) and 3T clinical scanners, its superior sensitivity makes 7T MRI a valuable tool to probe the brain’s subcortical gray matter at a much finer scale and with superior tissue contrast^24–26^. The increased signal-to-noise ratio provides unique opportunities to develop time-efficient imaging sequences for sub-millimeter quantification of changes specific to certain thalamic nuclei, and to regions which have never been studied in this context before due to difficulties to identify them at lower field strengths, such as the claustrum^25,26^. Here, we will extract several MRI-based (micro)structural features to detect subcortical anatomical changes using group-level comparisons, per structure and metric. In parallel, we will use a multivariate data-driven approach to investigate covariance patterns between the subcortical MRI phenotype (i.e., combining all MRI features) and clinical profiles as well as the importance of the different features towards these covariance patterns.

## Material and methods

### Study cohort

Participants were recruited from two prospective studies of patients with drug-resistant focal epilepsy: the EPINOV trial (NCT03643016) and the local EPI study. Both studies were approved by the local ethics committee, and all participants gave written consent. We retrospectively included all patients with a 7 Tesla (7T) brain MRI and a SEEG between June 2017 and October 2020. Fifty-two patients were initially included in this study. Five patients were excluded for suboptimal quality of MRI data and two patients were excluded because SEEG was performed before the MRI. A last patient was excluded because of a large cerebral pial angioma that altered the anatomy and prevented adequate brain segmentation. Finally, data from 44 patients were used for further analyses. Patients were classified into five groups according to the anatomo-functional organization of the EZN as defined by visual and quantitative SEEG data analyses^27^ by expert neurologists (authors F.B. and J.M.): (i) temporal plus, with EZN either limited to the temporal regions or with maximal epileptogenicity within the temporal lobe but EZ extension to some extratemporal regions (e.g., temporo-insular, temporo-frontal, temporo-occipital, N=25), (ii) prefrontal, with EZN involving prefrontal regions (N=8), (iii) posterior, with EZN involving parietal and/or occipital regions (N=5); (iv) insulo-opercular, with EZN involving the insula and/or opercular cortex (N=3) and (v) central-premotor, with EZN involving the primary motor and/or premotor cortex (N=3). A second grouping-level was added by separating between temporal (i.e., temporal plus group patients, ‘TLE’, N=25) and non-temporal patients (all other groups, ‘NTLE’, N=19) together to gain statistical power. Finally, we recruited 33 age- and sex-matched healthy controls for comparison.

### Image acquisition

All data were acquired on a Siemens Magnetom 7T MRI system (Siemens Healthineers, Erlangen, Germany) equipped with a 32-channel head coil (Nova Medical, Inc., Wilmington, MA USA). After an automatic B_0_ shimming procedure, a whole brain B_1_^+^ map was acquired using a spin-echo based sequence by assessing the ratio of consecutive spin and stimulated echoes (WIP#658, Siemens Healthineers). Then, for anatomical analyses, a whole brain three-dimensional (3D) Magnetization Prepared with 2 Rapid Acquisition Gradient Echoes sequence (MP2RAGE) was acquired^26^. MP2RAGE is an extension to the 3D MPRAGE sequence which combines two Gradient-Recalled Echo (GRE) volumes acquired at separate inversion times following an adiabatic inversion pulse. This sequence produces a T_1_-weighted (T_1_w) image with a spatially normalized contrast and reduced field bias. Acquisition parameters were: TR_mpr_/TR/TE/TI_1_/TI_2_ = 5000/7.4/3.13/2750 ms, α_1_/α_2_ = 6°/5°, FOV = 240 mm (matrix size: 402 × 402), 256 partitions, a parallel imaging acceleration factor of 3 (2D GRAPPA), and Partial Fourier factors of 6/8 in both phase- and partition-encoding directions. This resulted in a nominal isotropic voxel size = 0.6 mm and a total acquisition time of 10.12 min.

### Image preprocessing

The high resolution MP2RAGE data were used to segment thalamic nuclei and other basal ganglia for extracting morphometric and T_1_ features. Pre-processing of the MP2RAGE anatomical data included (i) correction for gradient non-linearities using the ‘*gradunwarp*’ tool provided by the Human Connectome Project (HCP, https://github.com/Washington-University/gradunwarp); (ii) post-hoc T_1_ correction for residual B_1_ ^+^ inhomogeneities; (iii) a skull stripping workflow optimized for MP2RAGE data using ‘PreSurfer’ and (iv) cortical surface reconstruction and parcellation using FreeSurfer (v7.1.1).

The MP2RAGE data were first corrected for B_1_^+^ inhomogeneities as described in the original paper resulting in a ‘corrected’ T_1_w (i.e., UNI) and quantitative T_1_ map, synthesized using a lookup table specific for the current sequence setup^28^. Non-brain tissue was then removed using the MP2RAGE’s second inversion and (corrected) UNI images using SPM12 functionalities wrapped within PreSurfer (https://github.com/srikash/presurfer)^29,30^. The skull stripped MP2RAGE T_1_w images were used as input for the sub-millimeter processing workflow implemented in FreeSurfer (v7.1.1, http://surfer.nmr.mgh.harvard.edu/) image analysis suite to obtain brain tissue segmentations, estimated total intracranial volume (eTIV) and white matter and pial surface reconstructions^31^.

### Segmentation of thalamic nuclei and basal ganglia

Isolation of brain structures was achieved in two-fold: (i) through an in-house developed thalamic nuclei (and basal ganglia) atlas (‘7TAMIBrain’)^32^ and (ii) by applying the Thalamus Optimized Multi Atlas Segmentation (‘THOMAS’) method^33^. Both methods rely on multi-atlas label fusion (MALF) which has proven superior compared to single-atlas registration methods^34^.

#### 7TAMIBrain

An improved version of the recently published 7TAMIBrain pipeline by Brun et al. was employed^32^. While initially based on a single template-to-subject registration, current version makes use of 60 atlases constructed from the 7TAMIBrain dataset^35^. This multi-atlas approach renders the segmentation results more robust and accurate, especially in cohorts with (potential) variable anatomies. The most important steps are highlighted below but see the Supplementary Methods for more details.

For each subject, its skull stripped T_1_ map was first coregistered to each of the 60 atlases using the quick three stages registration procedure (i.e., rigid + affine + deformable SyN atlas-to-subject registrations), as implemented in ANTs ‘*antsRegistrationSyNQuick*’ script^36,37^.

Fixed target (i.e., subject to segment) and moving (i.e., atlases) images were cropped (with matrix size = 192^3^) and centered on the DGN atlases pre-segmented via the single-atlas pipeline to optimize the computing time.

Second, the Joint Label Fusion algorithm and ANT’s ‘*antsJointLabelFusion2*’ script was used to merge the warped atlases into the individual subject’s space^34^. Label fusion reduces the impact of potential biases from the image registration while providing a good spatial consistency for the label boundaries. By default, the Joint Fusion method is only performed in voxels for which the atlas’ labeling consensus is lower than 80%, otherwise Majority Voting is used to determine label assignment. For this work, Joint Fusion was used for any voxel for which full consensus was not reached across all atlases by setting this threshold at 100%. This reduces potential recurrent biases in the segmentation of epileptic subjects with large anatomical differences from the healthy controls of 7TAMIBrain dataset.

#### THOMAS

To assess the impact of atlas choice on the thalamic results, thalamic nuclei were additionally segmented using THOMAS^33^. Since THOMAS is optimized to work with white matter-nulled (WMn) Magnetization Prepared with Rapid Acquisition Gradient Echoes (MPRAGE) images (i.e., enhanced thalamic-white matter contrast), MP2RAGE T_1_ maps were first white matter-nulled using a recently proposed thresholding method based on the inversion recovery signal equation^38^:

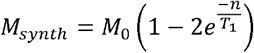

Here, a T_1_ threshold of 700 ms. was found to provide most reliable performance as assessed through visual inspection of segmentation results using varying thresholds. The WMn T_1_ maps were than used as input for the joint-fusion THOMAS pipeline (https://github.com/thalamicseg/thomas_new).

### MRI feature extraction

Multiple volume (size, T_1_ and deformation) and surface-based (vertex-wise displacement, area and curvature) features were extracted from the MRI data using the thalamic (both atlases) and basal ganglia segmentations (7TAMIBrain atlas only).

#### Volume-based features

For each ROI, total volume was computed (mm^3^) while longitudinal relaxation time (T_1_, ms) were averaged across voxels. In addition, voxel-wise deformations were obtained by warping the 7TAMIBrain T_1_ template to the subject’s T_1_ map (i.e., as part of the single-atlas procedure). Here, a non-linear registration was performed using ANTs’ deformable (diffeomorphic) registration (with shrink factors: 12×6×4×2×1; smoothing factors: 6×3×2×1×0 vox; max iterations: 100×100×70×50×10; transformation model: SyN; similarity metric: cross-correlation)^37^. The resulting affine transformation matrix and non-linear transformation field were combined to obtain a single displacement field. Local change in tissue density was then estimated as the derivative of the displacement of a given voxel in each direction. The derivatives can be calculated as the determinant of the Jacobian matrix using ANTs’ *ComputeJacobianDeterminant* function. The absolute value of the Jacobian determinant at voxel *p* gives us the factor by which the tissue expands or shrinks near *p*. By taking the logarithm of the determinant, no change is given by 0, tissue loss is given by a positive value, and tissue expansion is given by a negative value. Average deformation-based morphometry (DBM) values were obtained by averaging across all voxels within each ROI.

#### Surface-based features

To assess changes in shape, surface-based metrics were extracted following the procedure described previously and using openly available image processing scripts (https://github.com/khanlab/surfmorph)^39^. The subject’s T_1_w image and labels were first aligned along the AC-PC plane, following the HCP procedure, to enable left-right flipping to combine analyses of the left and right hemisphere labels. Fuzzy labels for each of the subject’s labels (ROIs) were then obtained by smoothing the binary ROI label with a 1 × 1 × 1 mm kernel size. The fuzzy (i.e., smoothed) labels for each of the ROIs, for each subject, were transformed to MNI space using linear transformations based on the subject’s MP2RAGE T_1_w to MNI volume transformation. These linearly aligned labels were used to generate unbiased averages for surface generation by iterating through steps of (i) template generation by averaging across subjects, and (ii) registration of each segmentation image to this template using LDDMM registration. The resulting fuzzy segmentation was then used to generate the ROI’s template surface through a 50% probability isosurface. The 3D volume of the ROI’s template was then fit to each subject’s segmentation using LDDMM, with affine initialization to provide vertex-wise correspondence between all surfaces of that specific ROI. The template surface was then propagated to each subject’s ROI, to provide surfaces with common indices for projecting vertex displacement values and extraction of additional shape features. As such, vertex area and surface curvature were extracted using the *‘surface-vertex-areas’* and *‘surface-curvature’* Connectome Workbench (v1.5.0) command line functions, respectively^40^. Please, see Supplementary Fig. 1 for a schematic of this procedure.

Please note that single surfaces were constructed for the thalamus (i.e., by grouping thalamic nuclei) and striatum (i.e., caudate nucleus, nucleus accumbens and putamen). However, nuclei-specific averages were calculated through thalamic and striatal surface atlases. To construct thalamic and striatal surface-based nuclei atlases, subject-specific volume labels were propagated towards the subject’s surface models along the closest vertex’ normal, after which Majority Voting (i.e., across all subjects) was used to assign a label to each vertex. ROI-wise displacement, area and curvature were then reduced to a single shape component through principal component analyses.

### Statistical analysis

Prior to statistical analyses, ROI-results were corrected for confounding, linear effects of age, head size (both continuous), and sex (male vs. female) and hemisphere (left vs. right) using the *confounds* (v.0.1.1) Python package^41^. Contributions of confounding variables to the given feature were determined using a linear regression model based on data obtained in control subjects only. Control and patient feature data were then ‘residualized’ by subtracting the contributions from the confound variables and z-scored using the controls’ mean and standard deviation. An overview of the impact of each of these confounding factors on the different metrics can be found in Supplementary Fig. 2. Eventually, a data matrix of N_subjects_ x [N_feature type_ x N_ROIs_ x N_sides_] was derived that was used as input for the following statistical analyses.

#### Feature-wise comparison

Assessment of statistical differences of MRI features between subject groups was performed using (multivariate) analyses of variance (ANOVA). Besides correcting for age, head size, sex and hemisphere (see paragraph on confounder correction), multivariate ANOVA (MANOVA) and/or Bonferroni multiple comparisons correction was used in case of across nuclei and/or pair-wise statistical comparisons, respectively. Moreover, for each feature, ipsilateral minus contralateral differences were computed to assess possible unilateral changes.

#### Partial Least Squares analyses

Partial Least Squares (PLS) analyses were carried out to further explore the variability in MRI-based features between groups (i.e., mean-centered PLS) and with respect to the clinical features across patients (i.e., behavioral PLS) using the *PyPLS* Python package (https://github.com/netneurolab/pypyls)^42,43^. Compared to mass-univariate (Pearson or Spearman) correlation analyses, PLS works well with data characterized by multicollinearity, as is the case for the current collection of MRI features. Mean-centered PLS is a multivariate statistical approach that defines sets of variables in a matrix (X_*n*×*p*_) which maximally discriminate between subgroups within the matrix. Here, the *n* rows represent the individual subjects and *p* columns the MRI-based features. The resulting latent variable (LV) then constitutes a weighted combination of all MRI-based features that best differentiates groups (e.g., ‘TLE vs NTLE vs. controls’). On the other hand, behavioral PLS identifies weighted patterns of variables across two given matrices (X_*n*×*p*_ and Y_*n*×*p*_) that maximally covary with each other. For example, when comparing patients’ MRI-based features in matrix X and patients’ clinical phenotype in matrix Y. Statistical evaluation of PLS results was performed using permutation tests (N=10000) to assess significance of overall patterns (i.e. LVs) and by bootstrap resampling (N=10000) to rank MRI and clinical features based on importance/reliability.

A graphical overview of the full analytical pipeline is displayed in Fig. 1.

**Figure 1.**
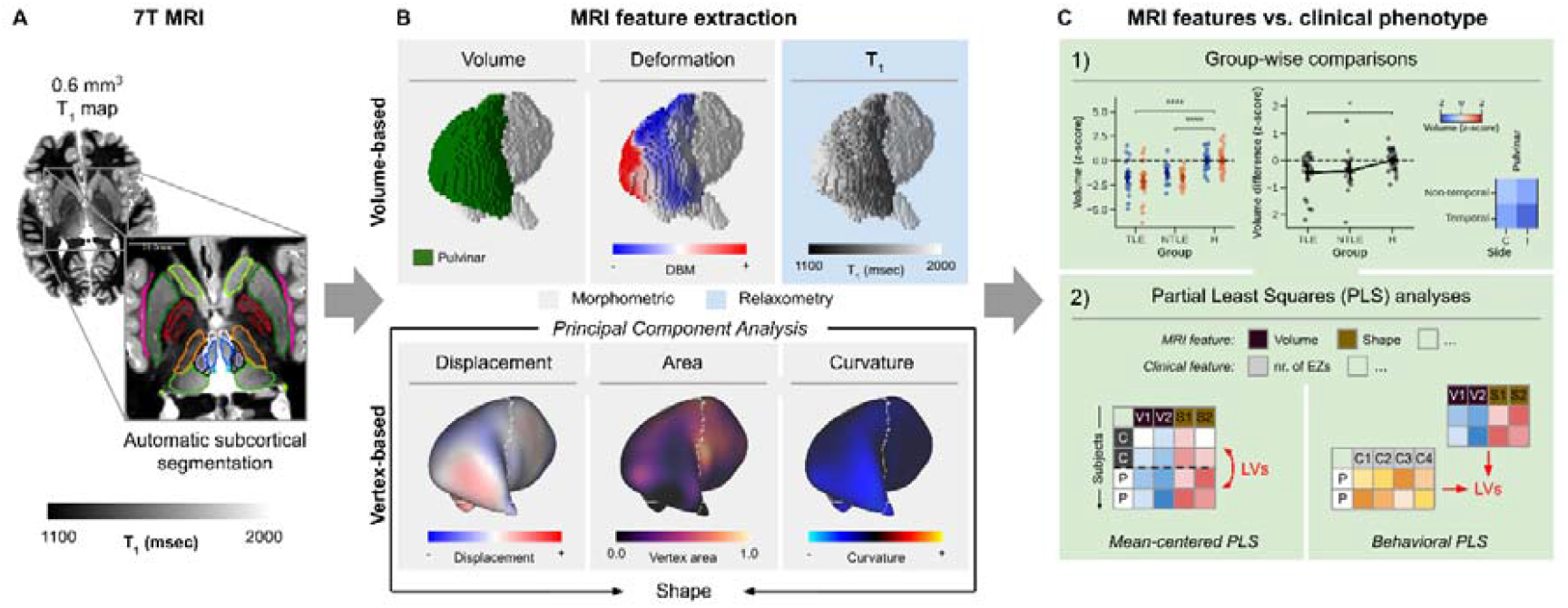
Schematic display of the analytical pipeline. (A) First, high resolution T_1_ maps were used to segment the thalamus and other basal ganglia. (B) Several MRI features were then extracted, focusing on morphometry and longitudinal relaxation time (T_1_). Surface-based metrics related to shape (i.e., displacement, area and curvature) were reduced to a single shape component. (C) Statistical analyses were performed for (1) group-wise comparisons as well as (2) identification of latent variables.

## Results

### Clinical features

Demographic characteristics and clinical data for all patients are shown in Table 1. The mean age at epilepsy onset was 12.6 (range 0.1–40) years, mean duration of epilepsy was 18.3 (range 2–47) years. Left and right seizure onset was observed in 23 (52%) and 21 (48%) cases, respectively, while on average 5.25 (range 1–14) regions were categorized as epileptogenic with a mean seizure frequency of 38.7 (range 2–300) per month. Anatomical MRI appeared normal in 25 (57%) and showed a structural abnormality in 19 (43%) of the cases based on 3T examinations. For one patient, the MRI showed a lesion at 7T while this was normal at 3T. Epilepsy etiology according to available histopathological findings included gliosis in seven, (suspected) focal cortical dysplasia (FCD) type I/II/III in respectively five, nine and one patient(s), dysembryoplastic neuroepithelial tumor (DNET) in four; polymicrogyria (PMG) in two and periventricular nodular heterotopia (PNH) in two patients.

**Table 1.** Patient clinical demographics.

| Demographic |  |  |  | Epilepsy |  |  |  |  |  |  | Lesion info |  |  |
| --- | --- | --- | --- | --- | --- | --- | --- | --- | --- | --- | --- | --- | --- |
| Patient | Age (yrs) | Sex | Group | Epilepsy type | Side | Age at onset (yrs) | Disease duration (yrs) | Seizure frequency (per month) | Frequency of bilateral tonic-clonic seizures (per year) | Number of epileptogenic regions | 3T MRI | 7T MRI | Etiology/<br>Histological findings |
| 1 | 21 | M | TLE | Temporal Plus | R | 0.8 | 20 | 10 | 4 | 12 | Lesion | Lesion | FCD2b |
| 5 | 22 | F | TLE | Temporal Plus | R | 13 | 9 | 150 | 0 | 8 | Normal | Normal | Unknown/no abnormality |
| 6 | 25 | F | TLE | Temporal Plus | L | 10 | 15 | 2 | 0 | 1 | Normal | Normal | FCD1 |
| 7 | 40 | M | NTLE | Prefrontal | R | 16 | 24 | 90 | 0 | 8 | Lesion | Lesion | Gliosis |
| 8 | 40 | M | NTLE | Posterior | R | 4 | 36 | 12 | 0 | 2 | Normal | Normal | Unknown/no abnormality |
| 12 | 56 | F | NTLE | Posterior | R | 30 | 26 | 20 | 0 | 7 | Lesion | Lesion | Suspected FCD2 |
| 13 | 44 | M | NTLE | Insulo-opercular | L | 13 | 31 | 2 | 0 | 3 | Normal | Normal | Unknown/NA |
| 16 | 46 | M | NTLE | Centra-Premotor | L | 4 | 42 | 120 | 0 | 3 | Lesion | Lesion | FCD2/NA |
| 18 | 48 | F | TLE | Temporal Plus | R | 40 | 9 | 30 | 0 | 3 | Normal | Normal | FCD1 |
| 20 | 33 | F | TLE | Temporal Plus | R | 24 | 9 | 4 | 0 | 3 | Normal | Normal | FCD3a |
| 22 | 29 | F | NTLE | Prefrontal | L | 12 | 17 | 20 | 0 | 2 | Lesion | Lesion | DNET |
| 24 | 27 | F | NTLE | Centra-Premotor | L | 4 | 23 | 90 | 0 | 4 | Lesion | Lesion | FCD2b |
| 25 | 37 | F | TLE | Temporal Plus | L | 12 | 25 | 16 | 0 | 1 | Lesion | Lesion | Ganglioglioma |
| 27 | 24 | F | TLE | Temporal Plus | L | 21 | 3 | 10 | 0 | 12 | Normal | Normal | Unknown/NA |
| 29 | 28 | F | NTLE | Prefrontal | L | 0.1 | 28 | 90 | 0 | 2 | Normal | Lesion | FCD2a |
| 30 | 36 | M | NTLE | Posterior | R | 3 | 33 | 300 | 0 | 3 | Lesion | Lesion | FCD2b |
| 31 | 21 | F | NTLE | Insulo-opercular | R | 14 | 7 | 60 | 0 | 1 | Lesion | Lesion | DNET |
| 35 | 31 | F | NTLE | Prefrontal | L | 8 | 22 | 30 | 0 | 2 | Lesion | Lesion | FCD2b |
| 37 | 47 | F | TLE | Temporal Plus | L | 5 | 42 | 10 | 0 | 6 | Normal | Normal | Unknown/NA |
| 38 | 29 | M | TLE | Temporal Plus | L | 3 | 26 | 120 | 0 | 2 | Normal | Normal | Mild gliosis |
| 39 | 25 | M | TLE | Temporal Plus | L | 11 | 14 | 8 | 4 | 6 | Normal | Normal | Unknown/NA |
| 40 | 23 | M | TLE | Temporal Plus | L | 12 | 11 | 8 | 0 | 4 | Normal | Normal | FCD1 |
| 45 | 26 | M | NTLE | Prefrontal | L | 17 | 10 | 50 | 0 | 6 | Normal | Normal | FCD2a |
| 53 | 22 | F | TLE | Temporal Plus | R | 4 | 18 | 100 | 0 | 11 | Normal | Normal | Unknown/NA |
| 54 | 15 | M | TLE | Temporal Plus | L | 2 | 13 | 5 | 0 | 7 | Normal | Normal | FCD2a |
| 55 | 57 | F | TLE | Temporal Plus | R | 35 | 22 | 5 | 6 | 4 | Lesion | Lesion | Gliosis |
| 56 | 22 | M | NTLE | Posterior | L | 20 | 2 | 9 | 0 | 2 | Lesion | Lesion | PMG |
| 57 | 17 | M | NTLE | Insulo-opercular | L | 7 | 10 | 12 | 1 | 5 | Normal | Normal | Unknown/NA |
| 58 | 52 | F | TLE | Temporal Plus | L | 5 | 47 | 12 | 0 | 7 | Lesion | Lesion | PMG, PNH |
| 60 | 30 | M | TLE | Temporal Plus | L | 3 | 26 | 15 | 6 | 5 | Normal | Normal | Unknown/NA |
| 61 | 27 | F | TLE | Temporal Plus | R | 17 | 10 | 5 | 1 | 6 | Normal | Normal | Unknown/NA |
| 65 | 42 | F | TLE | Temporal Plus | L | 28 | 14 | 15 | 6 | 4 | Lesion | Lesion | Gliosis |
| 66 | 36 | M | TLE | Temporal Plus | R | 29 | 7 | 60 | 12 | 3 | Normal | Normal | FCD1 |
| 67 | 41 | M | NTLE | Prefrontal | R | 12 | 29 | 60 | 0 | 2 | Normal | Normal | Unknown/NA |
| 70 | 18 | F | TLE | Temporal Plus | R | 13 | 5 | 15 | 6 | 6 | Normal | Normal | Mild gliosis |
| 71 | 38 | F | TLE | Temporal Plus | R | 20 | 18 | 68 | 0 | 4 | Lesion | Lesion | PNH, FCD1 |
| 74 | 30 | F | TLE | Temporal Plus | L | 19 | 21 | 2 | 0 | 6 | Normal | Normal | Unknown/NA |
| 76 | 20 | M | NTLE | Prefrontal | R | 7 | 13 | 15 | 0 | 11 | Lesion | Lesion | Pediatric infiltrative glioma |
| 78 | 31 | M | TLE | Temporal Plus | R | 22 | 9 | 4 | 0 | 5 | Normal | Normal | Mild gliosis |
| 80 | 21 | M | TLE | Temporal Plus | L | 12 | 9 | 2 | 0 | 3 | Lesion | Lesion | Ganglioglioma |
| 81 | 16 | M | NTLE | Posterior | R | 10 | 6 | 10 | 12 | 14 | Normal | Normal | Unknown/NA |
| 82 | 21 | F | NTLE | Prefrontal | R | 9 | 11 | 5 | 0 | 1 | Normal | Normal | Mild gliosis |
| 83 | 16 | M | TLE | Temporal Plus | R | 1,5 | 14 | 30 | 0 | 13 | Lesion | Lesion | DNET |
| 84 | 17 | M | NTLE | Centra-Premotor | L | 0.5 | 17 | 2 | 0 | 11 | Lesion | Lesion | DNET |

Abbreviations: M/F, male/female; TLE, temporal lobe epilepsy; NTLE non-temporal lobe epilepsy; L/R, left/right; FCD, focal cortical dysplasia; NA, not applicable; DNET, dysembryoplastic neuroepithelial tumor; PMG, polymicrogyria; PNH, periventricular nodular heterotopia.

### MRI features

Figure 2 shows the full data matrix with rows representing subjects, grouped based on epilepsy type, and columns representing the individual MRI features organized per metric type (i.e., volume, deformation, shape and T_1_), brain structure (thalamic nuclei and other basal ganglia) and side with respect to seizure onset for patients (ipsilateral vs contralateral), while based on hemisphere for control subjects. From this birds-eye view, both volume and T_1_ show lower z-scores (i.e., brighter blue) for the patients compared to controls while the opposite pattern is true for deformation and shape (i.e., brighter red).

**Figure 2.**
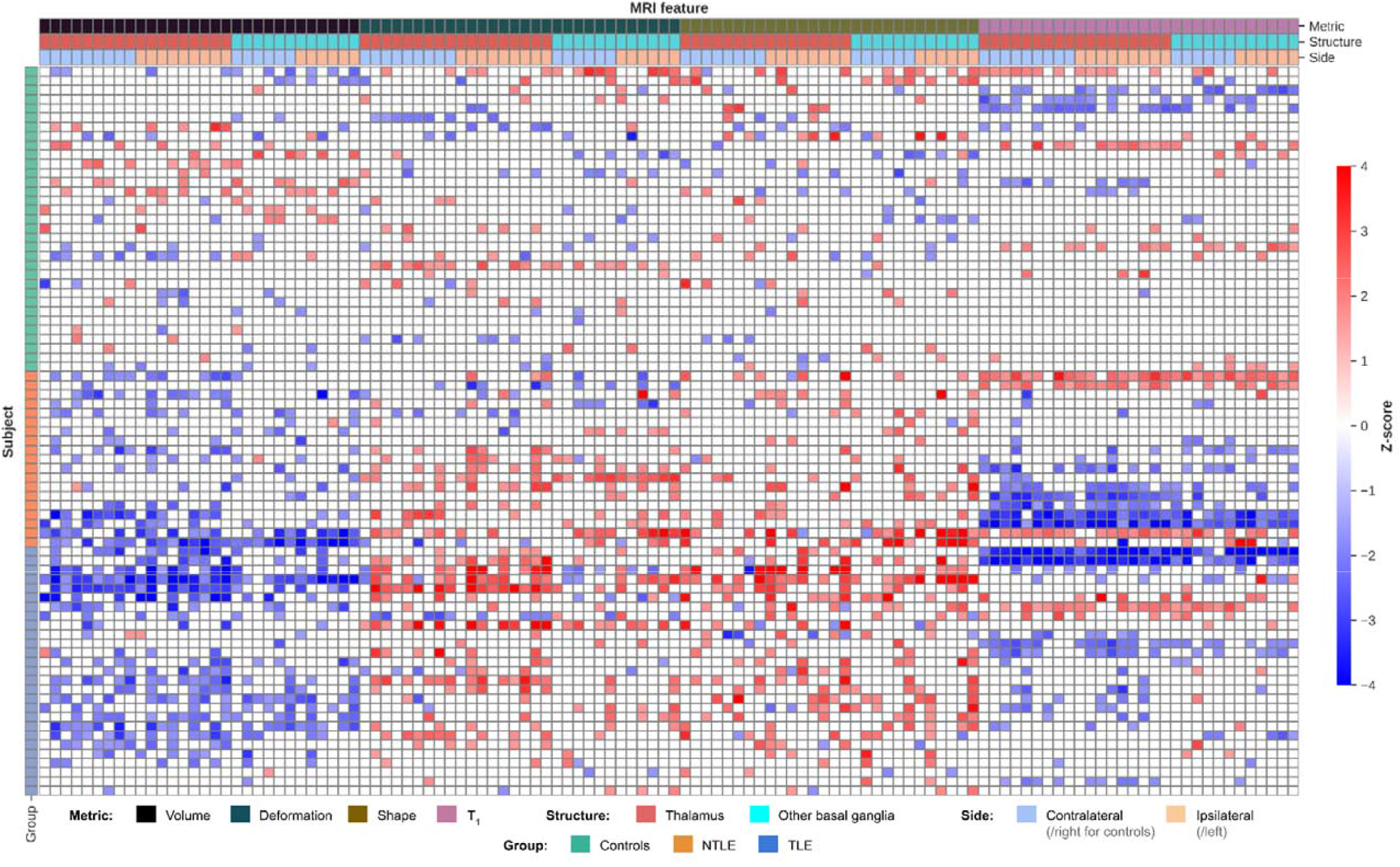
Z-scores heatmap. Individual MRI features are color-coded (thresholded at -1.5 and 1.5) as indicated by the horizontal bars on the top according to metric type (volume, deformation, shape and T_1_), brain structure (thalamus vs. other basal ganglia) and side with respect to seizure onset for patients (contra-vs. ipsilateral), while left vs right for control subjects. Vertical color bars on the left side indicate subject group.

In the following sections will present these results in more detail focusing first on thalamic volume and T_1_, after which these will be analyzed in conjunction with the remaining MRI features (deformation and shape), other basal ganglia data, as well as patient-specific clinical features using PLS analyses.

#### Volume

Previous work has shown thalamic volume reduction in TLE. To confirm these effects in the current study cohort, as well as to investigate potential TLE and NTLE and nuclei-specific differences, we compared thalamic changes at a global scale (i.e., whole thalamus) as well as at the single nucleus-level.

At the level of the whole thalamus (Fig. 3A), ipsilateral and contralateral thalamic volumes were significantly different between groups (F_2,76_ = 20.68, p < .001). Both groups of patients had lower volumes compared to controls (p < .001). Thalamic volume was lower in the thalamus ipsilateral to the epileptogenic zone compared to controls in TLE (p < .05), but not in NTLE (p = .06). No significant differences were observed between TLE and NTLE.

**Figure 3.**
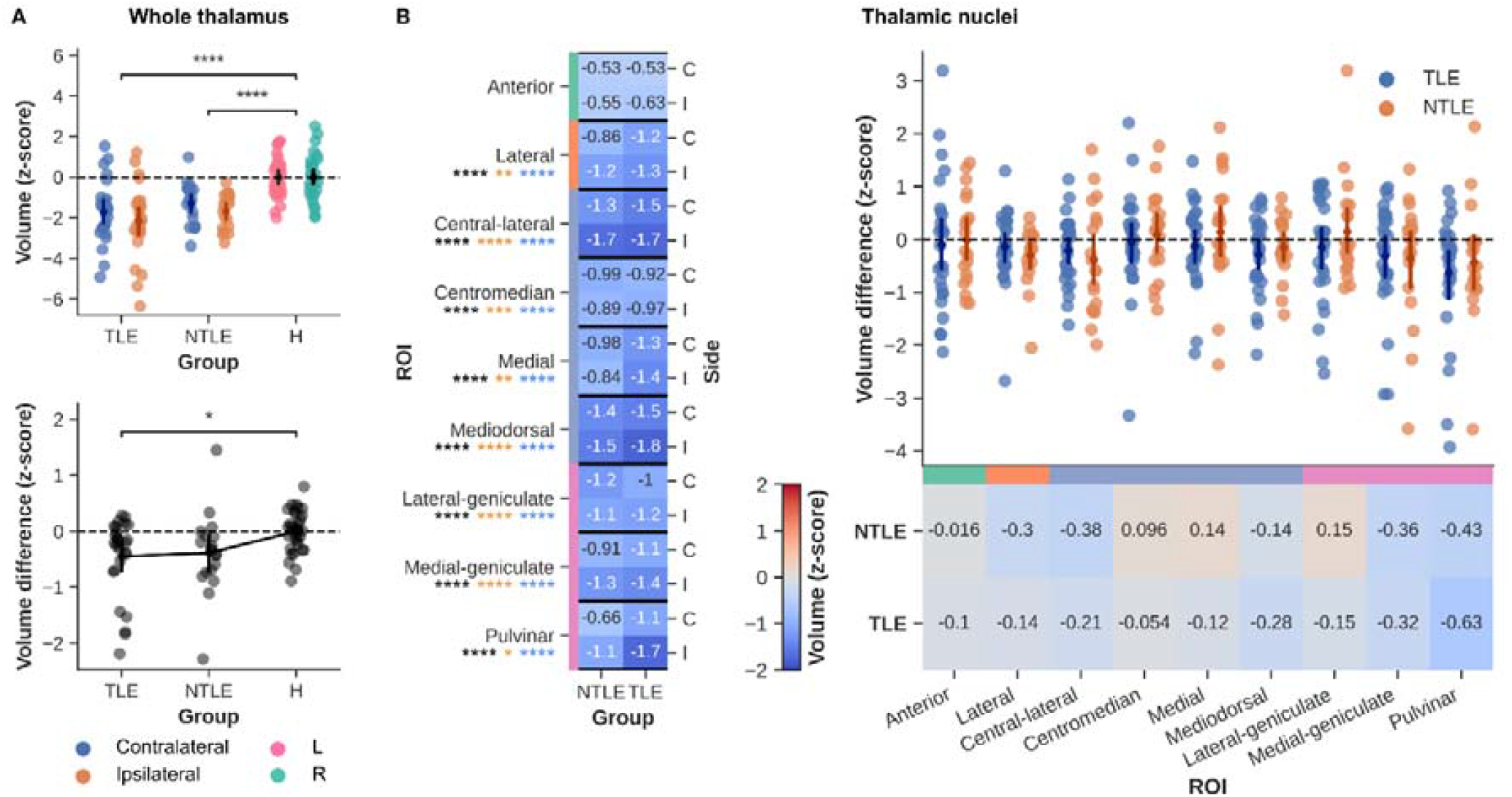
Volumetric results. (A) Whole thalamic volume (z-scored) was compared between groups (top panel) as well as between sides (i.e., ipsilateral-contralateral differences for patients, left-right for controls, bottom panel). (B) Similarly, average volumes of each thalamic nuclei (z-scored) are shown for both patient groups (columns) and side (rows, left heatmap). Asterisks below nuclei labels indicate MANOVA (black), and pair-wise comparison (orange for NTLE and blue for TLE, vs. controls) results. Strip plot and right heatmap in B illustrate the ipsilateral-contralateral T_1_ differences for each patient (color-coded by group). Color-bars alongside nuclei labels indicate group assignment. C = contralateral, I = ipsilateral, * p < .05, ** p < .01, *** p < .005, **** p < .001, Bonferroni corrected in case of multiple comparisons.

Thalamic nuclei volumes (Fig. 3B) were lower than controls in both focal epileptic groups (F_18, 138_ = 3.502, p < .001; Wilk’s Λ = 0.471, partial η^2^ = .31). The strongest general group effect was observed in the mediodorsal nucleus (F_2,77_ = 22.038, p < .001) with differences originating from both TLE and NTLE (both p < .001) groups compared to controls. The weakest (and only not significant) group effect was observed for the anterior nucleus (F_2,77_ = 2.956, p = .058). Asterisks below nuclei labels indicate MANOVA (black), and pairwise comparison (orange for TLE and blue for NTLE, vs. controls) results. Multivariate analysis across thalamic nuclei did not show systematic ipsi-vs. contralateral volume differences between groups.

For brevity reasons, see the Supplementary Results (section ‘Deformation and shape’) for extension of these analyses to the other morphometric features (deformation and shape, respectively).

#### T_1_ relaxometry

Thalamic T_1_ relaxometry values (Fig. 4A) were significantly modulated by group (F_2,76_ = 3.80, p < .05), with differences coming from the TLE patients compared to controls (p < .05).

**Figure 4.**
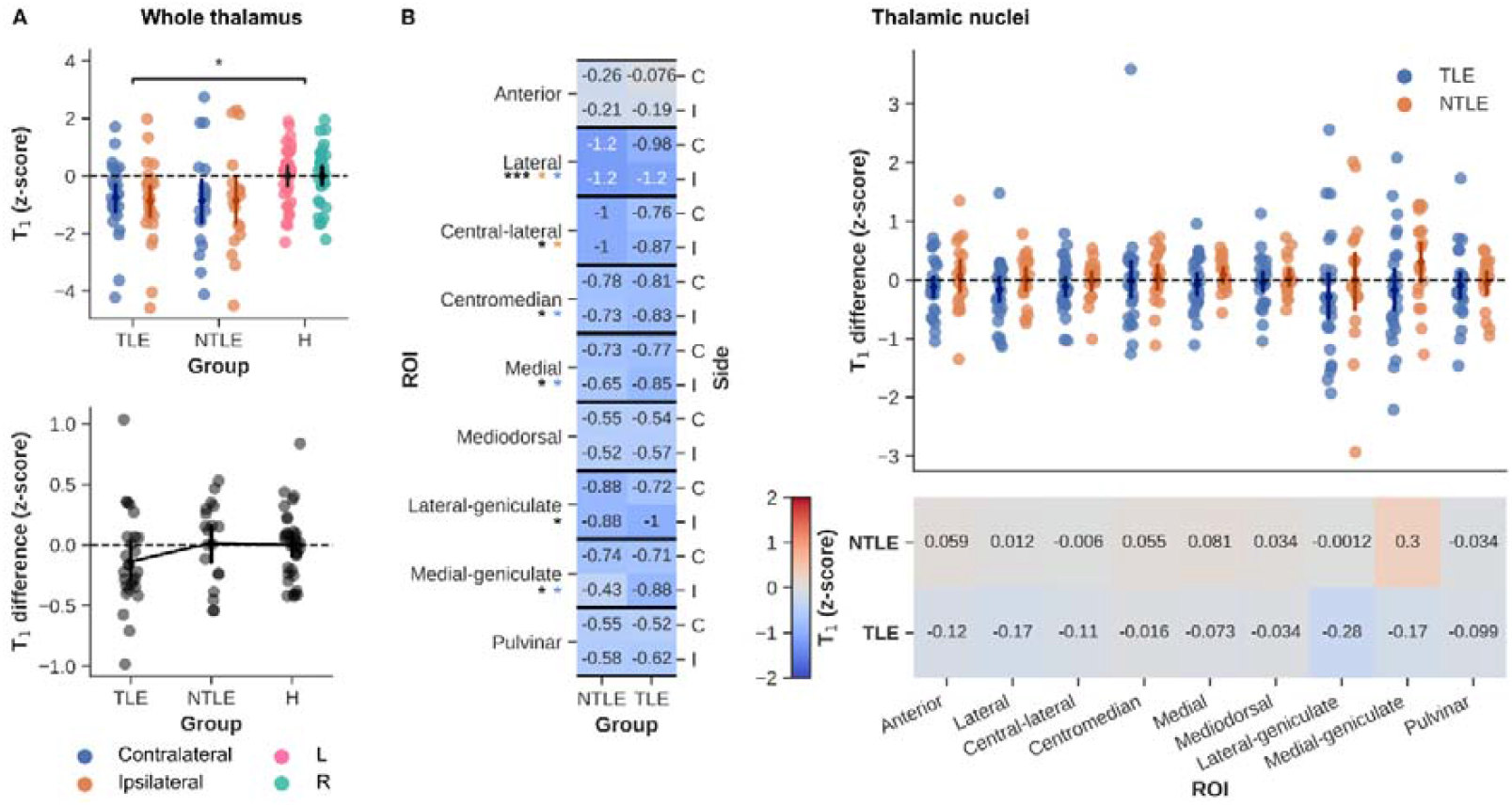
T_1_ results. (A) Whole thalamic T_1_ (z-scored) was compared between groups (top panel) as well as between sides (i.e., ipsilateral-contralateral differences for patients, left-right for controls, bottom panel). (B) Similarly, average T_1_ of each thalamic nuclei (z-scored) are shown for both patient groups (columns) and side (rows, left heatmap). Asterisks below nuclei labels indicate MANOVA (black), and pairwise comparison (orange for NTLE and blue for TLE, vs. controls) results. Strip plot and right heatmap in B illustrate the ipsilateral-contralateral T_1_ differences for each patient (color-coded by group). Color-bars alongside nuclei labels indicate group assignment. C = contralateral, I = ipsilateral, * p < .05, ** p < .01, *** p < .005, **** p < .001, Bonferroni corrected in case of multiple comparisons.

As for volume, thalamic nuclei T_1_ (Fig. 4B) values were lower than controls in both focal epileptic groups (F_18, 13)_ = 2.230, p < .01; Wilk’s Λ = 0.600, partial η^2^ = .24). Here, the lateral group shows the strongest effect (F_2,77_ = 3.228, p < .005) with similar contributions from TLE (p < .05) and NTLE (p < .05) patients. As for volume, the weakest effect was seen for the anterior nucleus (N.S.). T_1_ was not significantly different between the ipsi- and contralateral sides.

### MRI vs. clinical features

In previous sections we reported on thalamic changes, focusing on the MRI features separately. Nonetheless, it remains unclear which feature(s) provide(s) the most discriminatory power between groups and/or across patients when analyzed together as well as the added value of the other basal ganglia. As such, in the following we performed mean-centered and behavioral PLS analyses to rank MRI and clinical features in terms of importance/reliability, respectively.

#### Mean-centered PLS

Mean-centered PLS allows to rank individual MRI features based on their contribution to LVs that significantly differentiate(s) between subject groups. Contrasting MRI features between NTLE, TLE and controls resulted in one significant LV (81.4% explained variance, p_perm_ < .001). Fig. 5A shows the ranking and comparison of the MRI features (color-coded as in Fig. 2) in terms of their absolute contribution to the LV as defined by their bootstrapped ratio (BSR) between the feature’s loading and its 95% confidence interval. Metric, structure, and side-wise averages are shown using black diamonds (± 95% CI). Here, BSRs were significantly modulated by metric type (F_3,102_ = 12.27, p < .001) with higher BSRs observed for the volumetric features compared to deformation-, shape- and T_1_-based features (all p < .001). Although different in magnitude, a clear spatial correlation between volume and shape features can be observed when projecting the BSR value onto their respective surface reconstructions, with higher BSR values towards the ipsilateral, posterior thalamus (Supplementary Fig. 3).

**Figure 5.**
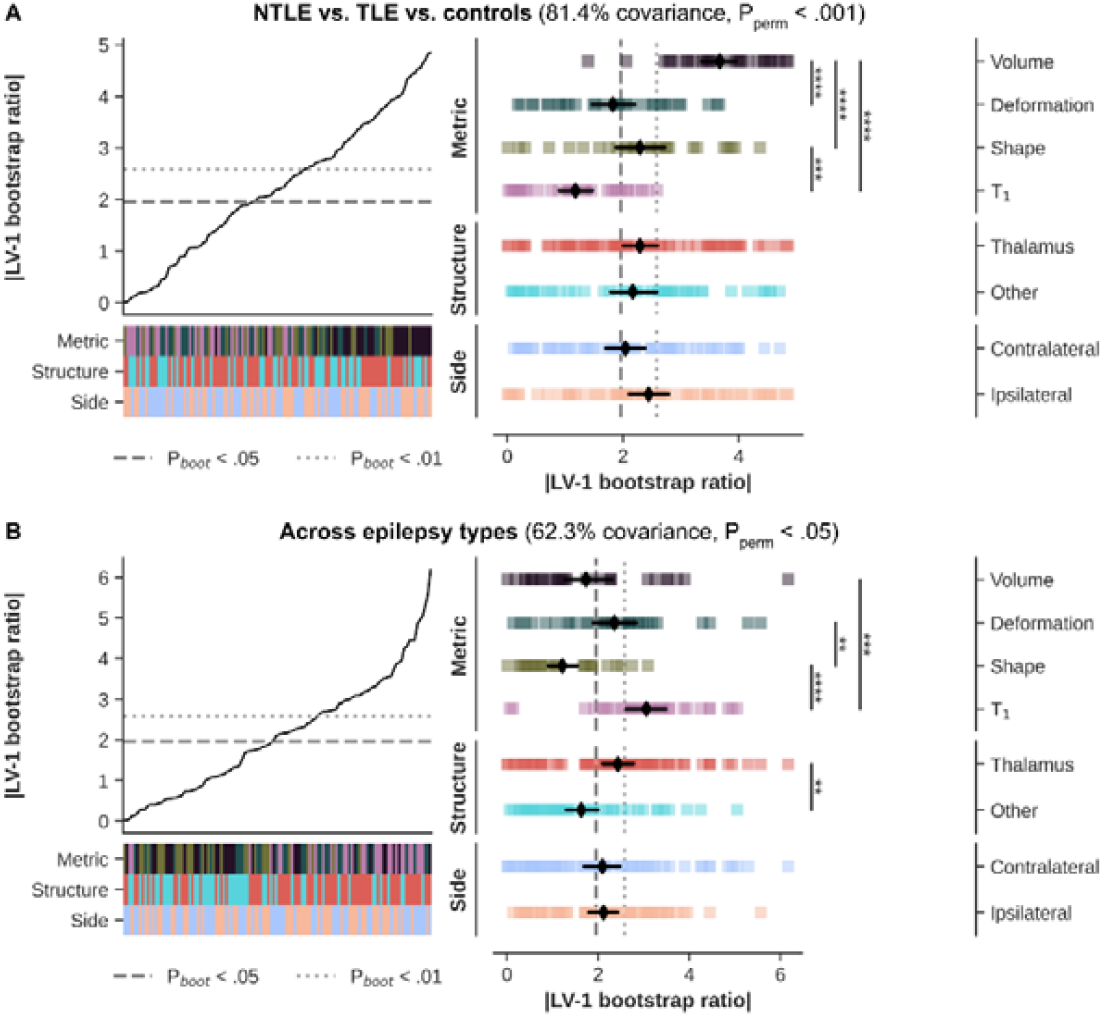
Mean-centered PLS results. (A) Line plot shows sorted bootstrapped ratios when contrasting NTLE, TLE and controls. Heatmaps indicate feature properties (metric type, structure, and side, color-coded as in Fig. 2). Strip plot shows the decomposition of features based on feature properties with black diamonds indicating average BSR (± 95% CI). (B) Same as A but contrasting epilepsy types (i.e., temporal plus, prefrontal, posterior, insulo-opercular and central-premotor). * p < .05, ** p < .01, *** p < .005, **** p < .001, Bonferroni corrected in case of multiple comparisons.

A weaker, but nonetheless significant LV was detected when contrasting the five epilepsy types (62.3% explained variance, p_perm_ < .05). Again, BSRs varied as function of metric type (F_3,102_ = 12.27, p < .001), but structure as well (F_1,102_ = 11.71, p < .005). Here, highest BSRs were observed for the T_1_-based features (p < .005 compared to volume and p < .001 compared to shape). Moreover, changes in thalamic nuclei appear more strong contributors than changes in the other basal ganglia (p < .005). See Supplementary Fig. 4 for the projection onto the surfaces.

#### Behavioral PLS

Focusing on the patient data only, behavioral PLS provides a mean to evaluate patterns based on both MRI and clinical features, independent of epilepsy type. This revealed one significant LV (35.2% explained variance, p_perm_ < .05). Here BSRs were significantly modulated by metric type (F_3,102_ = 13.91, p < .001), structure (F_1,102_ = 6.814, p < .05) and side (F_1,102_ = 7.680, p < .01). Contributions predominantly came from T_1_ (p < .001 compared to deformation-based features), thalamic (p < .05 compared to other basal ganglia) and contralateral (p < .05 compared to ipsilateral) features (Fig. 6A). With regards to the clinical characteristics, strongest contributions were found for the number of epileptic zones, but also presence of MRI-detectable lesion(s), disease duration (yrs) and seizure frequency (all p_boot_ < .05, Fig. 6B). Pink bars indicate associations that were significant at p_boot_□<□0.05. Surface projections reveal that volume features contributing robustly to the LV were found predominately within the thalamus, while T_1_ features were found more widespread across the basal ganglia (Supplementary Fig. 5). Again, volume and shape show similar surface-wise projections.

**Figure 6.**
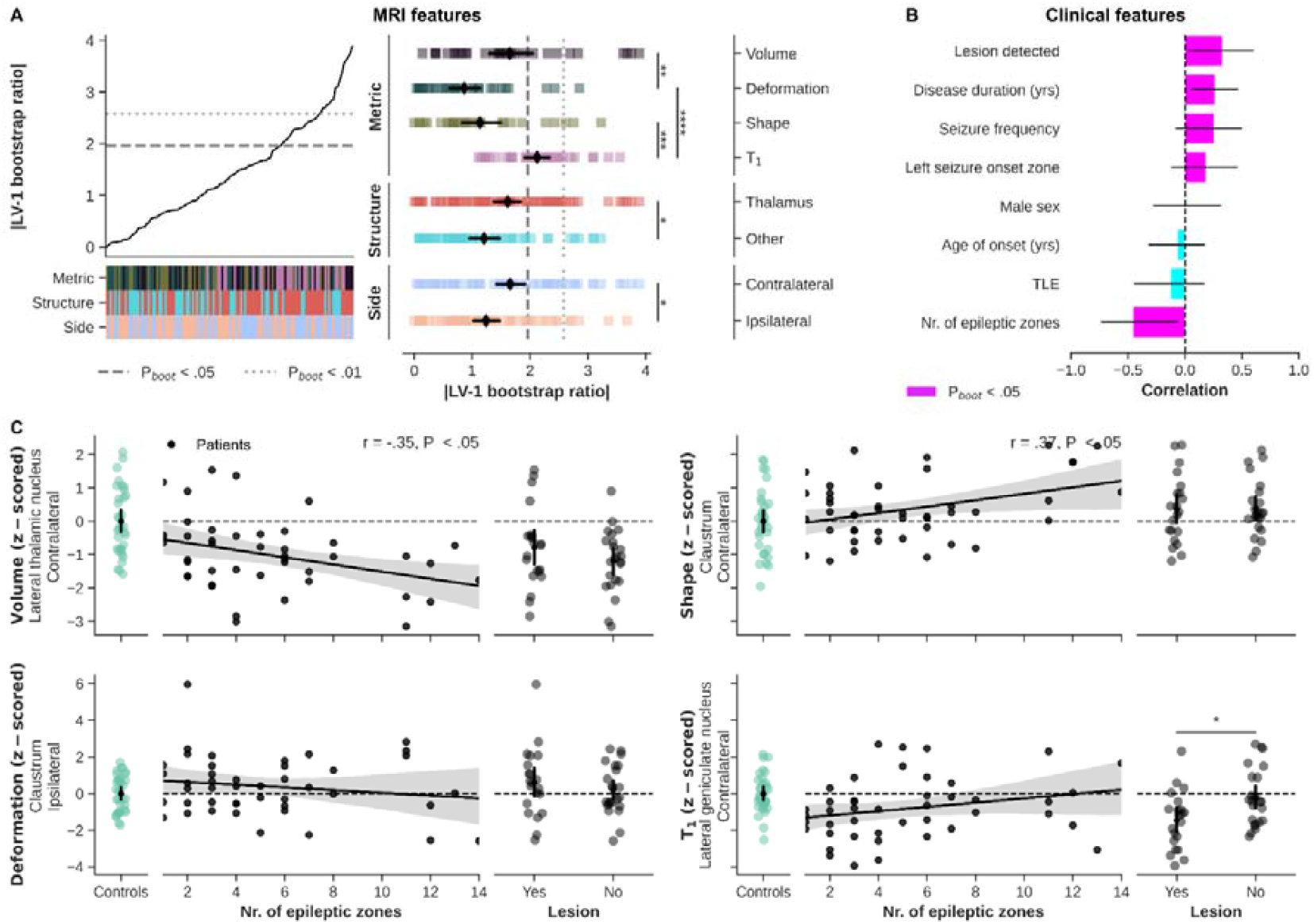
Behavioral PLS results (A) Line plot shows sorted bootstrapped ratios with heatmaps indicating feature properties (metric type, structure, and side, color-coded as in Fig. 2). Strip plot shows the decomposition of features based on feature properties with black diamonds indicating average BSR (± 95% CI). (B) Ranking of clinical features in terms of correlation with the LV. (C) Correlation between clinical features and MRI-based z-scores. * p < .05, ** p < .01, *** p < .005, **** p < .001, Bonferroni corrected in case of multiple comparisons.

Finally, the scatterplots in Fig. 6C visualize the correlations between covarying MRI and clinical features (number of epileptic zones and visible lesion, yes/no). We choose to show the MRI features with the highest BSR for each metric type. Here, lateral thalamic volume was lower in patients characterized by an increased number of epileptic zones, r(42) = -.35, p < .05. A similar (but inverse) pattern is visible for the claustrum’s shape, r(42) = .37, p < .05). T_1_ in the lateral geniculate nucleus seem to be reduced in patients with a visible lesion compared to patients without a lesion (p < .05).

### Replicating using THOMAS

To assess the dependence of current results on the atlas choice, all analyses were repeated but using the THOMAS atlas for the thalamic segmentations. These can be found in the Supplementary Results (section ‘Replicating using THOMAS’). Briefly, analyses conducted using thalamic results using the THOMAS atlas showed the same direction of differences as with the 7TAMIBrain atlas. Nonetheless, while remaining below significance threshold using the 7TAMIBrain atlas, the impact of seizure onset side was stronger across the different metrics using THOMAS. For example, average ipsi-vs. contralateral T_1_ differed significantly between TLE and NTLE patients (p < .05).

## Discussion

In this study we used 7T MRI anatomical data to characterize subcortical gray matter alterations in focal epilepsy patients. Our cohort of focal epilepsy patients showed structural abnormalities across the thalamus and basal ganglia for an array of morphometric metrics, as well as T_1_ as a measure of tissue microstructure. In general, the strongest impact was observed for TLE patients with the magnitude of inter-group differences varying based on the structure of interest. Moreover, the spatial pattern of these differences and the contribution of the different metrics varied depending on whether the patient data was contrasted against controls or among themselves.

### Global thalamic changes

Focal epilepsy patients showed significant reduction in thalamic volume, as well as increased – although characterized by smaller effect sizes – tissue deformation and shape values. These indicators of atrophy are in line with prior research in TLE^20,21,44^ and as recently demonstrated on a larger scale using the multisite ENIGMA Epilepsy dataset^10^. Here, we confirm this phenomenon in our TLE and NTLE focal epilepsy cohort. This is not surprising given the widespread thalamo-cortical connectivity^45^ and its implications in seizure initiation, propagation and spread^14,18,46^. Moreover, the ipsilateral predominance of global thalamic atrophy could intuitively be explained by the thalamo-cortical connectivity during seizures. This has been illustrated previously by accurate prediction of seizure onset laterality in TLE patients based on individual thalamo-hippocampal functional connectivity scores^47^.

Most interestingly, we are the first to report longitudinal relaxation data for the basal ganglia in both TLE and NTLE drug-resistant patients characterized by SEEG. In contrast to the data reported in the cortex^13^, we found a decrease in thalamic T_1_. It is known that T_1_ shortens with increasing myelin and iron (i.e., ferritin) concentrations^48^. Therefore, the observed decrease in T_1_ can be attributed to higher levels of myelin and/or iron in patients compared to controls. In particular the latter is a hallmark of several neurological and neurodegenerative diseases^49^. In epilepsy, using pilocarpine-treated rats to induce a status epilepticus, iron (and calcium) deposits were selectively observed across the thalamus^50^. Potential mechanisms leading to iron accumulation include increased blood–brain barrier (BBB) permeability, inflammation, iron redistribution within the brain, and changes in iron homoeostasis^49^. It remains unclear, however, if these abnormalities are mainly caused by seizures, or by the evolution of the disease and neurodegenerative processes. Leek et al (2021) showed that volumetric abnormalities of structures known to be important for the generation and maintenance of focal seizures are established at the time of epilepsy diagnosis and are not necessarily a result of the chronicity of the disorder^51^. On the other hand, longitudinal tracking of cortical thickness in TLE patients showed progressive thinning with longer disease duration, particularly in regions structurally connected with the ipsilateral hippocampus^52^.

Furthermore, differences between cortical and thalamus tissue and their increase^13^ vs. decrease in T_1_, respectively, might be explained based on their tissue’s organization including their connectivity. Cortical cyto- and myeloarchitecture is organized in a laminar fashion along a sheet-like structure with cortical myelination gradually decreasing towards the pial surface and layer-specific feed-forward and feedback projections^53^. This laminar organizational principle is not in place for thalamic gray matter with nuclei-specific (sub)cortical projections^54^. This imposes a dense network of innervating fiber bundles with high levels of crossing fibers^55,56^. Given the important role of the thalamus in seizure organization and therefore widespread participation in epileptiform activities, changes in thalamic T_1_ might be dominated by pathophysiological processes (e.g., BBB dysfunctioning^57^) that cause iron to accumulate^14,49^. In contrast, demyelination as also observed in postsurgical tissue derived from cortical dysplasia might be the predominant driver of the increase in cortical T_1_^58^. Also, while atrophy has been observed across both cortical and subcortical areas, differences in the microstructural environment (e.g., macromolecular composition) might render the impact of neuronal loss on T_1_ relaxation differently^59,60^. Nonetheless, based on T_1_ relaxation time alone, it remains difficult to explain the histological and molecular translation of the observed changes. Multivariate analyses with other quantitative metrics, like the tissue’s transverse relaxation time (T_2_*), susceptibility value (QSM) and diffusion properties (e.g., fractional anisotropy and mean diffusivity) are necessary to better explain this observed decrease in T_1_. This has proven to be beneficial at the cortical level with better discrimination of normal and abnormal neuron density in neocortical gray matter when T_1_ is combined with FA^61^, leading to better separation between left and right TLE^62^.

### Multi-scale structural alterations

The thalamus as a whole and by itself provides relevant information to better understand the disease processes in focal epilepsy. However, the human thalamus hosts several nuclei with varying cytoarchitectonic features and projections innervating distinct (sub)cortical regions^55,63^. Therefore, thalamic nuclei might be differentially affected depending on EZ (i.e., epilepsy type), epileptic network characteristics and other clinical features. Moreover, thalamic connectivity with basal ganglia warrant investigation of covarying patterns among subcortical structures^64–66^. Precise delineation and quantification of thalamic nuclei and basal ganglia have become feasible with the advent of 7T MRI and the possibility to map T_1_ both at a submillimeter resolution and (relatively) fast using the MP2RAGE sequence^26^. As a result, multiple protocols have been proposed, including the one developed in-house^32^, to segment thalamic nuclei and basal ganglia from such data^33,38^. When we compared results from our in-house 7TAMIBrain atlas method with those from another popular approach (THOMAS), thalamic volumetric measurements appear relatively similar despite differences in segmentation algorithm and nuclei definitions. In addition, brain tissue segmentations based on (quantitative) MRI data like transmit field (B ^+^)-corrected MP2RAGE maps are known to be more robust and accurate than those based on uncorrected images affected by residual B_1_^+^ biases^28,67,68^. In parallel, current results provide a proof-of-concept to assess morphometric and T_1_ properties of the claustrum – a deep brain structure characterized by challenging anatomy with its thin sheet of neurons, enclosed by white matter and situated between the insula and the putamen^69^ – in a clinical population.

Using mean-centered and behavioral PLS^42,43^, we then performed multivariate mapping of MRI-based subcortical features and patient characteristics (i.e., epilepsy type or clinical features) to isolate latent clinical-anatomical dimensions of focal epilepsy. Understanding the heterogeneity of clinical and anatomical manifestations of focal epilepsy remains a major challenge^10^. Multivariate analyses have been used across multiple neurological and neuropsychiatric disorders that are characterized by a high heterogeneity across patients and provided insight on how disease processes of brain structure and function are intertwined^43,70,71^. When applied on our multi-scale subcortical MRI phenotypes, containing morphometric and microstructural information for each structure and subject, mean-centered PLS analyses identified a prominent LV that explained 81.4% of variance between TLE, NTLE and control groups. Across patients only, hereby stratifying the phenotypes based on epilepsy type, the principal LV captured 62.3% of variance between groups while joint processing of the patients’ MRI and clinical phenotypes resulted in a LV that described 35.3% of their covariance. Permutation tests revealed that statistical significance was reached for each of these LVs while a bootstrapping procedure allowed us to rank the list of MRI and clinical features based on importance^42^. A closer inspection and comparison of these LVs revealed several interesting findings. First, while volume-based measures appeared to be most discriminative when control data were included, T_1_ measurements gain importance with respect to other metric types when discriminating across patients only. This was evident based on the MRI features only (i.e., mean-centered PLS), as well as when combined with the clinical features (behavioral PLS). Second, the spatial patterns of the LVs appear to be characterized by different topological modes with robust features either (i) confined to a set of thalamic nuclei or (ii) spread across the entire subcortex. When focusing on the thalamus again, morphometric abnormalities within the posteromedial thalamus, including the pulvinar and mediodorsal nucleus, acted as dominant discriminators between patients and controls. On the other hand, epilepsy types were best separated by their T_1_ within the posterolateral portion of the thalamus, in particular. Together, these patterns follow the same trend as the group-level analyses and suggest that predisposition of thalamic nuclei to alterations^50^ may depend on the property being evaluated but also on clinical characteristics, such as the location and/or number of epileptogenic zones. The medial pulvinar has been implicated in focal epilepsy based on frequently observed ictal activity changes during temporal lobe seizures^18,19^ and its atrophy^20^, and has been proposed as potential target for therapeutic deep brain stimulation (DBS) in drug-resistant epilepsies^72^. Similarly, given its connectivity with the amygdala and temporal cortex^56^, the mediodorsal nucleus has been studied in the context of (medial) temporal lobe epilepsy. For example, rats experiencing status epilepticus and then developing epilepsy had a decrease in volume of the mediodorsal thalamic region related to histological indicators of neuronal loss^73^. Extending these findings to humans, mediodorsal and posterior thalamic gray matter density was reduced in TLE patients^74^ and in patients with persistent seizures compared to patients without seizures after surgery^75^. On the other hand, spatial specificity of T_1_ decreases towards the lateral thalamus in focal epilepsy (as well as across epilepsy types) could be explained by the interaction between activity-regulated myelination^76^ and seizure pathophysiology. Increased seizure network myelination after epilepsy onset has recently been demonstrated in rat models of absence seizures and generalized epilepsy^77^. This, together with the strong connectivity of the lateral thalamus to somatosensory, posterior and parietal areas, organized along an anterior-posterior axis, might explain the consistent decrease in T_1_ for the lateral thalamic nuclei in our patient cohort^56^. The anterior nucleus volume and T_1_ appeared relatively unaffected. This is surprising given the frequent involvement of the limbic system in the TLE group. Potential explanations could be its small size, shape variability (as also shown in this work)^78^, and its close vicinity to WM (i.e., internal medullary lamina) which can influence segmentation accuracy. Furthermore, anterior nucleus atrophy is particularly associated with hippocampal sclerosis, which was not present in our cohort^79^.

However, in line with earlier work using structural covariance networks in TLE and idiopathic generalized epilepsy^10^, our LVs show that volume-based patterns extend beyond the thalamic borders and involve basal ganglia such as internal globus pallidus and striatum as well. This pattern might represent the same shared patterns of structural abnormalities across different epilepsies as suggested by Whelan et al. (2018), despite the heterogeneity of epilepsy and seizure types. Our results complement other recent studies of multivariate analyses in focal epilepsy. For example, Lee et al. (2022)^71^ identified several MRI-based components spanning cortical and white matter tissue that described interindividual variability in TLE. Within a framework to predict treatment response or surgery outcome, these outperformed classifiers that did not operate on latent factor information. Another multivariate approach emphasized a close coupling of cognitive dysfunction and large-scale network anomalies in TLE^80^. Here we show that subcortical volume measures allow discrimination between patients and controls, while T_1_ measures look promising to further differentiate patients based on epilepsy type. Altogether, these multivariate results demonstrate the brain-phenotypic complexity of focal epilepsy at the whole-brain scale with the extent, interplay and type of changes depending on brain compartment as well as clinical phenotype.

### Methodological considerations and study limitations

Although the consistency across thalamic segmentation protocols corroborates the robustness of our results, they might not be generalizable to external datasets and those acquired at lower field strengths, in particular.

T_1_ is dependent on magnetic field strength with the level of dependency varying based on tissue characteristics (i.e., low vs. high molecular mobility)^81^. It has been shown that 7T allows detection of epileptogenic lesions not visible at 3T^23^. Therefore, based on typical acquisitions performed at 3T and 7T, differences in the signal- and contrast-to-noise ratios of T_1_ data will impact sensitivity to segment individual nuclei as well as the power to detect patient-control morphometric and T_1_ differences at lower field strengths^25,82^.

Inherent to the demographics of focal epilepsy^83^, we are limited in the number of patients diagnosed with extra-temporal epilepsies. Consequently, we have pooled all NTLE patients into a single patient group, with the risk of losing sensitivity during group-level comparisons. Moreover, the statistical significance of the PLS LVs were evaluated within a descriptive framework to detect above-chance associations in the current dataset. As such, we did not test whether such associations generalize to new data. Replication using a separate cohort with a different number of patients per epilepsy type might rank MRI features differently and lead to different conclusions. Evaluation of out-of-sample correlation and predictive performance using an independent and larger study cohort is needed. It is therefore our hope that in the future a similar action will be undertaken to establish a multisite 7T epilepsy data as achieved by the ENIGMA Epilepsy study group^84^. As a first step, recent efforts by the 7T Epilepsy Task Force led to practical recommendations on targeted use of 7T MRI in the clinical management of patients with epilepsy^23^. Such datasets will help mitigate problems related to limited sample sizes as well as means to investigate potential methodological biases.

### Conclusion and future directions

This work reveals multi-scale – based on multiple parameters across different structures – subcortical alterations in focal epilepsy patients using 7T MRI data. It shows that the relative importance for different MRI features across subcortical structures depend on whether the goal is to isolate patients from controls or to differentiate between epilepsy types. Moreover, these results provide a comprehensive representation of the subcortical phenotype of focal epilepsy and its correlation with clinical parameters. In future work, integration of (i) other quantitative MRI, as well as (ii) electrophysiological data might allow us to better understand its direct relation to the structures’ microstructural and epileptogenic properties, respectively. Moreover, extending this framework to the cortex provides opportunities to investigate the impact of subcortical alterations on the brain’s system-wide functioning. Finally, since DBS is largely based on empirical protocols, comprehensive mapping of subcortical changes and relationship with disease heterogeneity might contribute to an improved treatment planning and outcome.

## Supporting information

Supplementary Results

## Data and code availability

All code and notebooks to reproduce this work can be found online (https://github.com/royhaast/smk-epilepsy-7T-sctx). The data are not publicly available due to sensitive information that could compromise the privacy of research participants. Nonetheless, the data are available from the corresponding author on reasonable request.

## Declaration of Competing Interest

The authors declare no conflict of interest.

## Acknowledgments

We would like to thank the patients and control participants who agreed to take part in this study. The authors would like to thank Patrick Viout, Lauriane Pini, Claire Costes, and Veronique Gimenez for data acquisition and study logistics, and Lucas Gauer and Olivier Girard for their fruitful discussion on the presented results and/or support at various stages of this project. The study was financially supported by the French government under the “Programme Investissements d’Avenir”, Excellence Initiative of Aix-Marseille University -A*MIDEX (AMX-19IET-004), 7TEAMS Chair, EPINOV (ANR-17-RHUS-0004) and the European Union’s Horizon 2020 Framework Program (785907 and 945539).

## Supplementary Figures

**Supplementary Figure 1.**
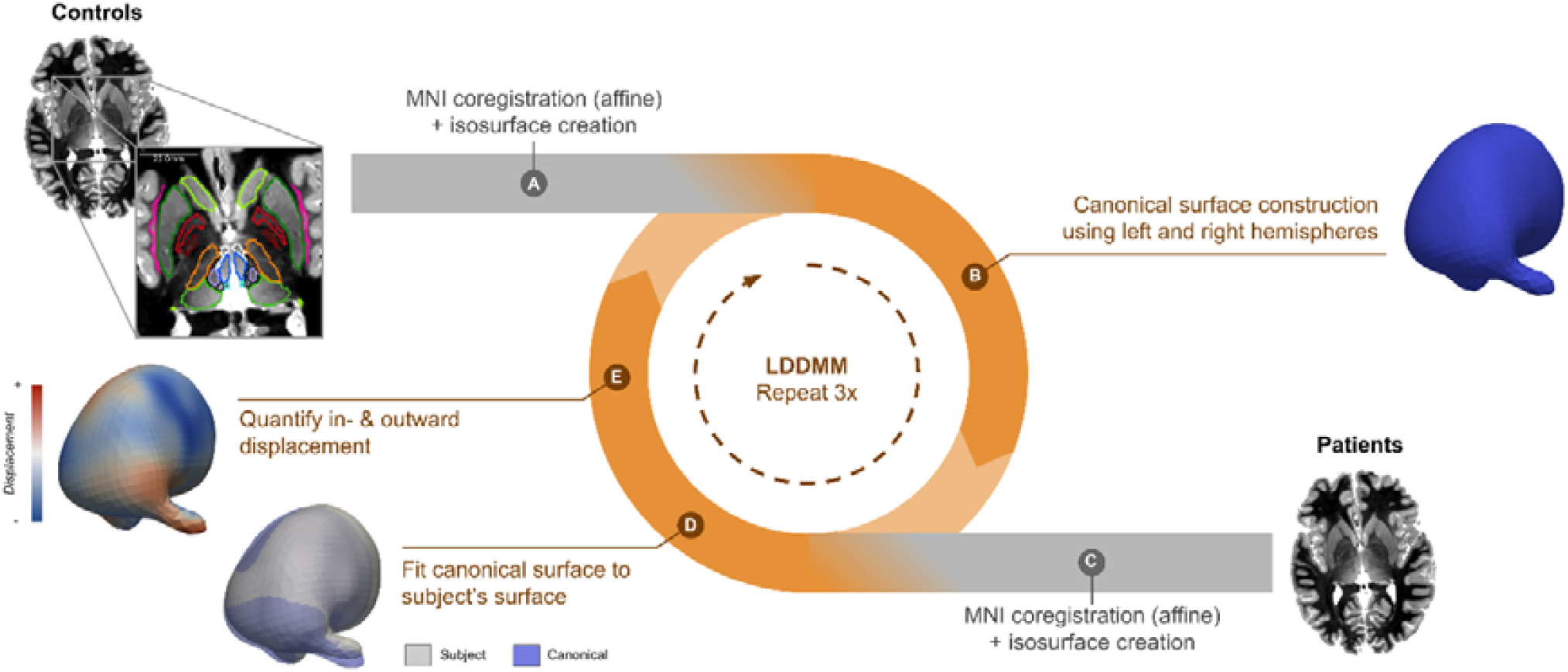
Schematic display of the surface-based morphometry pipeline. (A and C) Subject-specific subcortical surfaces in MNI space were first reconstructed from the segmentation results. (B) Surfaces from control subjects were then used to build a canonical surface that served as reference for vertex-wise quantification of in- and outward displacement through an iterative process (in orange, D-E).

**Supplementary Figure 2.**
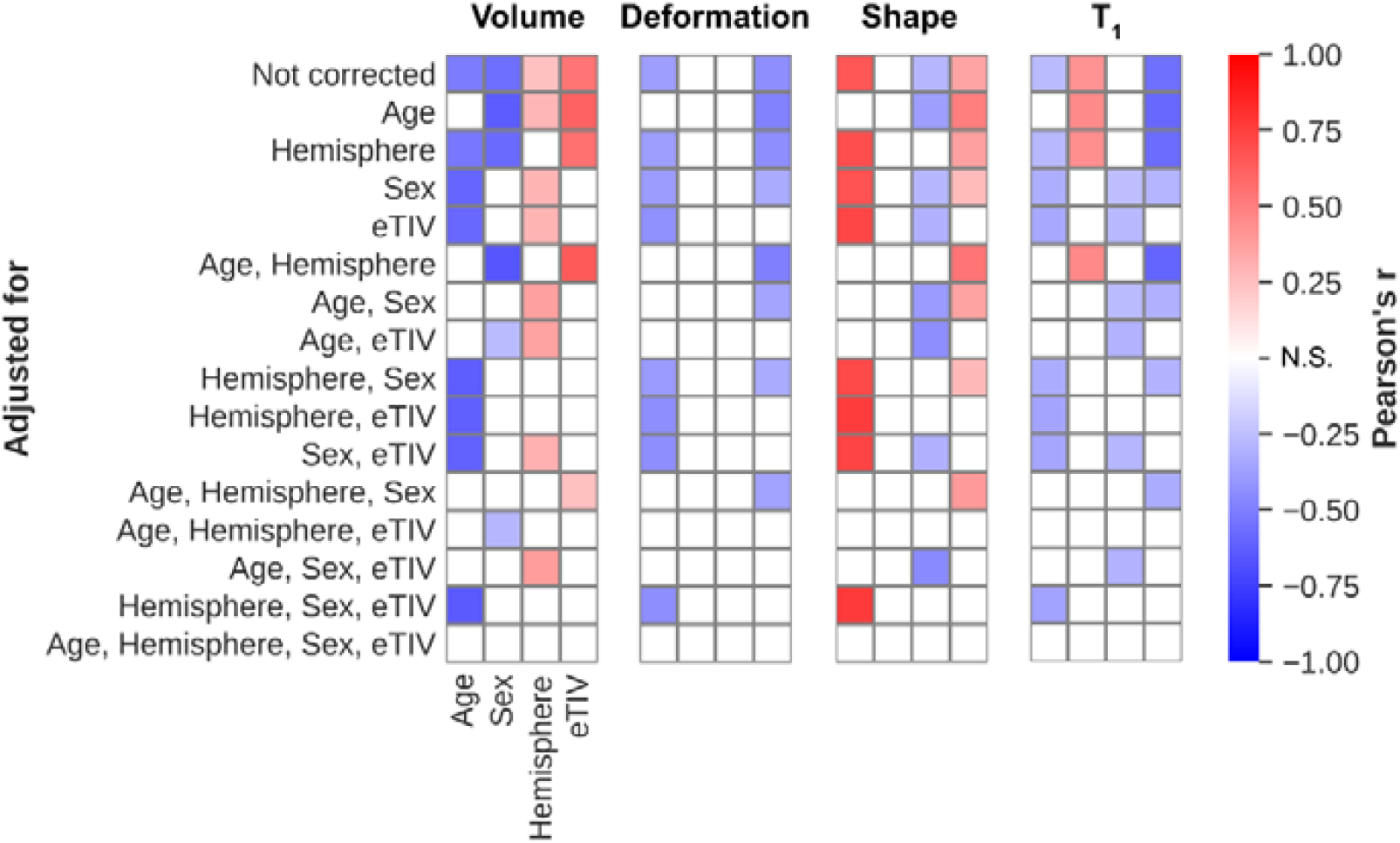
Impact of confounding factors (columns) on the different MRI metrics, with and without prior adjustment (rows). Significant positive and negative Pearson’s correlations between confounding values and MRI metrics are shown in red and blue, respectively.

**Supplementary Figure 3.**
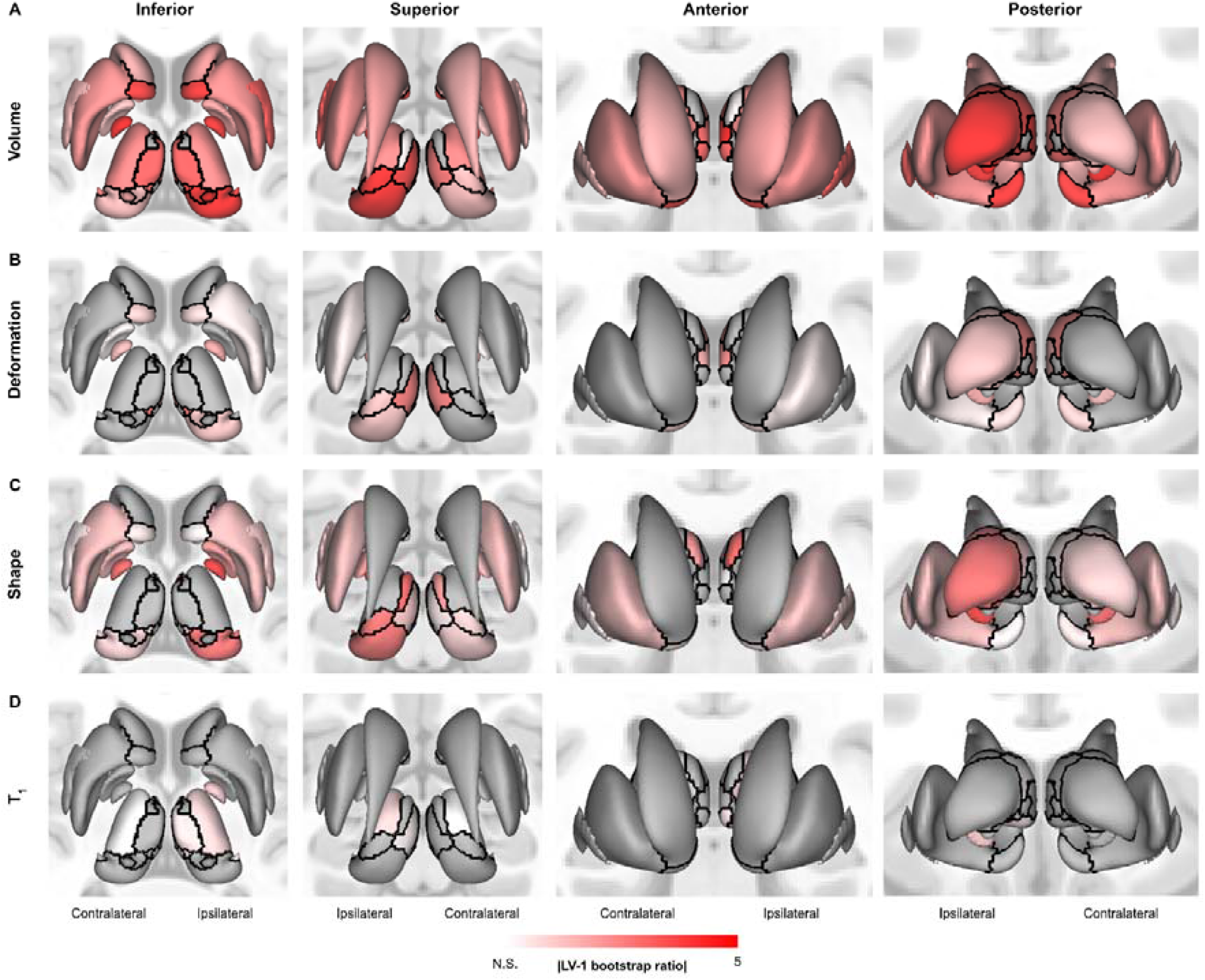
Projection of significant bootstrapped ratio values (color-coded), based on mean-centered partial-least squares analyses (i.e., temporal lobe vs non-temporal lobe vs. controls), onto the structures’ surface reconstructions. Results are separated per MRI metric type (rows). Black solid lines indicate structure boundaries.

**Supplementary Figure 4.**
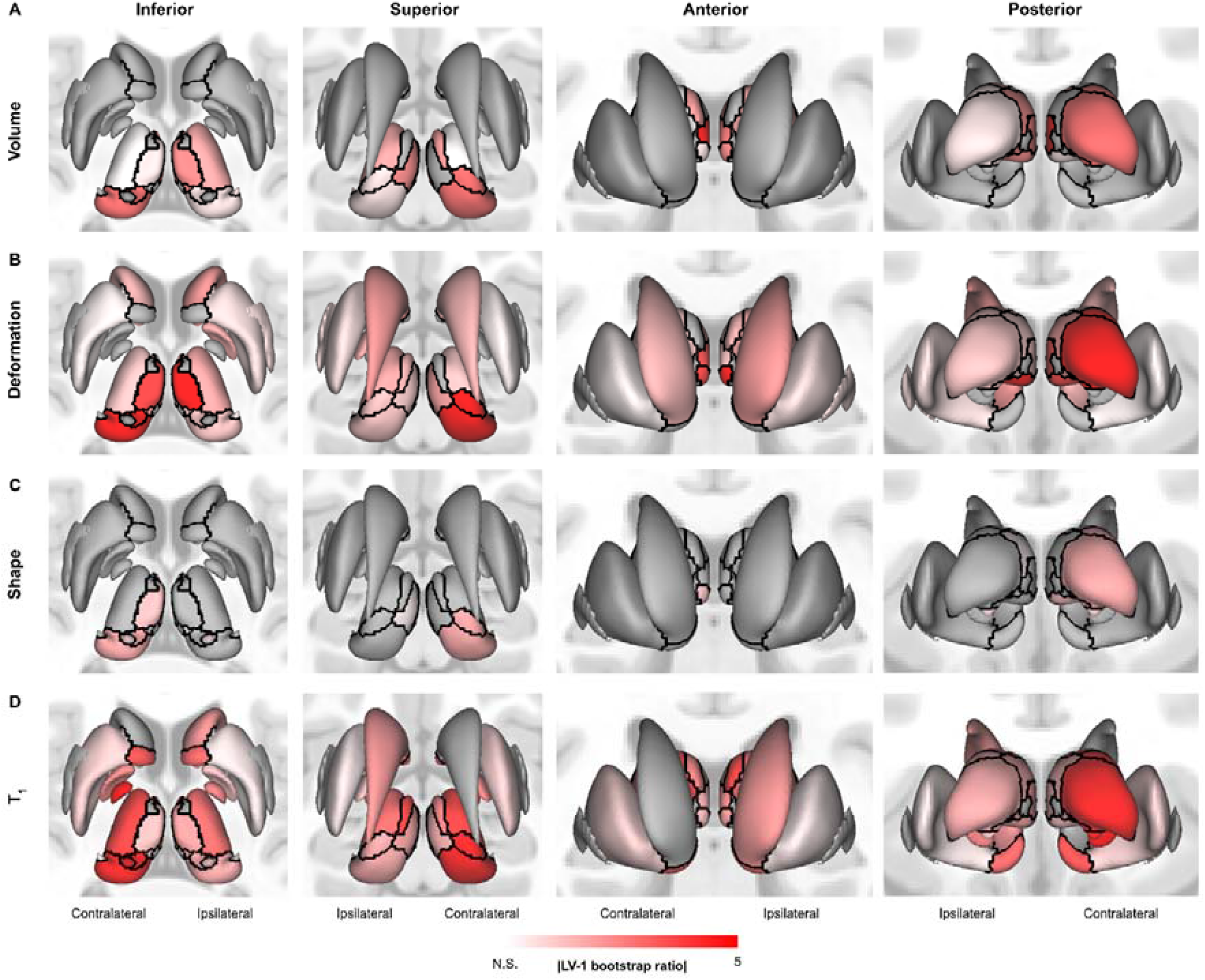
Projection of significant bootstrapped ratio values (color-coded), based on mean-centered partial-least squares analyses (i.e., between epilepsy types), onto the structures’ surface reconstructions. Results are separated per MRI metric type (rows). Black solid lines indicate structure boundaries.

**Supplementary Figure 5.**
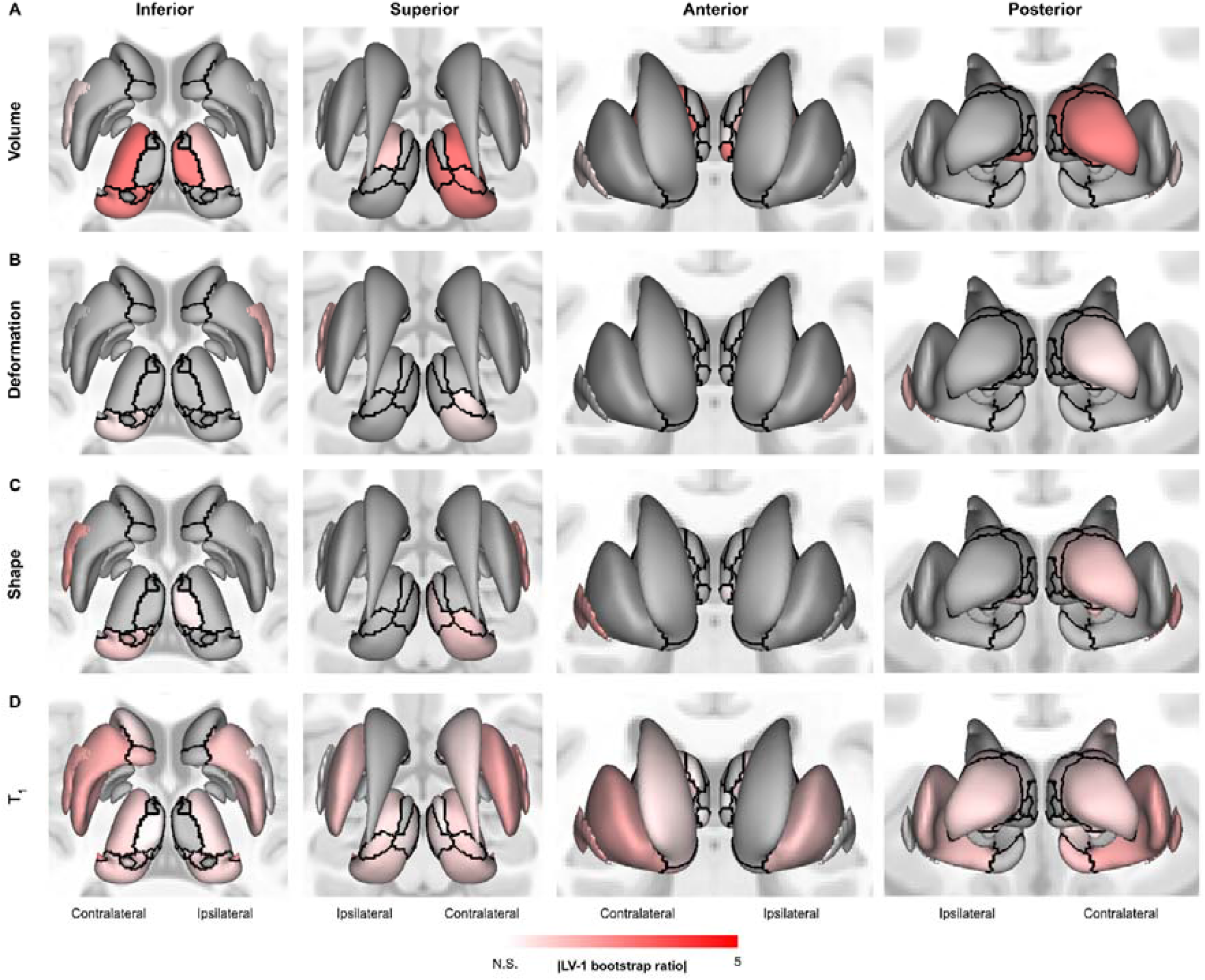
Projection of significant bootstrapped ratio values (color-coded), based on behavioral partial-least squares analyses (i.e., across patients using clinical characteristics), onto the structures’ surface reconstructions. Results are separated per MRI metric type (rows). Black solid lines indicate structure boundaries.

## Notes

### Competing Interest Statement

The authors have declared no competing interest.

### Summary of Updates

Updated manuscript to correspond with submitted version to journal. Slight change in title and added missing supplementary figures.

