## Supplementary Methods for "Multi-scale structural alterations of the thalamus and basal ganglia in focal epilepsy as demonstrated by 7T MRI"

### Multi-atlas label fusion using the 7TAMIBrain atlas

The full pipeline multi-atlas label fusion (MALF) pipeline using the 7TAMIBrain atlas is depicted in Figure M1.

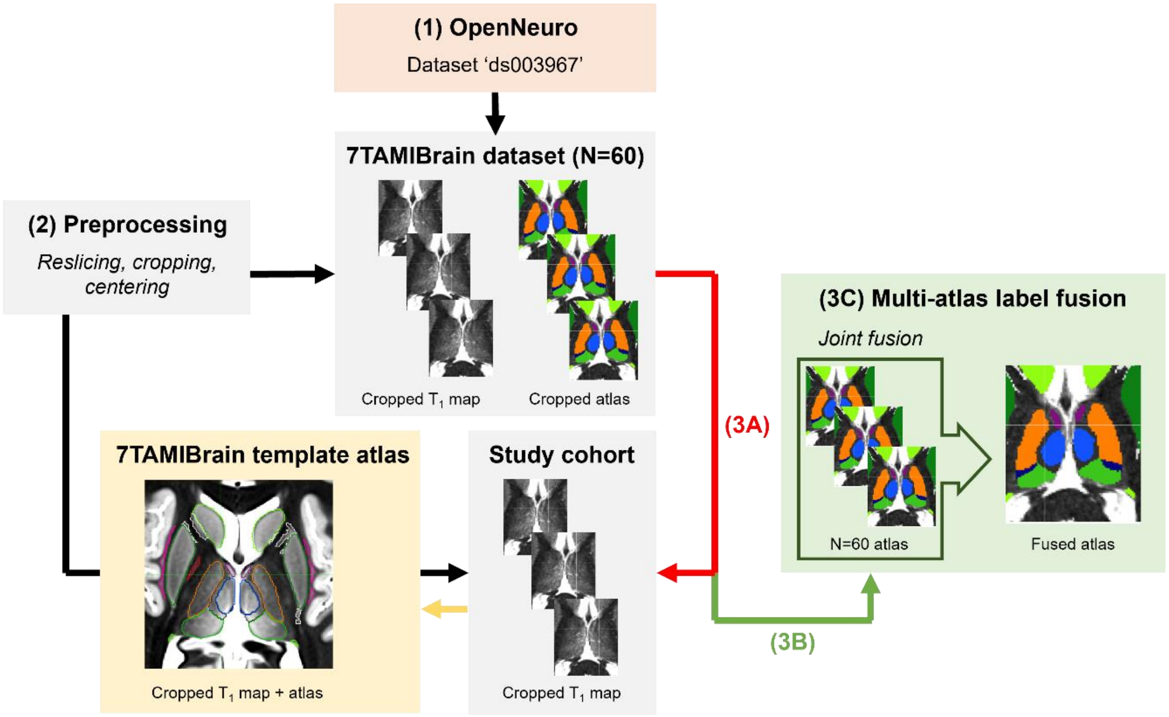


*Figure M1 – Schematic display of the full multi-atlas application based on the 7TAMIBrain atlas.*

**The multi-atlas pipeline stages:**

1. A set of N=60 $T_1$ maps and corresponding subcortical segmentations, forthwith referred to as the 'atlas database', were obtained through the 7TAMIBrain dataset (identifier 'ds003967') in the OpenNeuro repository[1].
2. To optimize the computing time as well as registration efficiency during the subsequent steps, input images were first centered and cropped around the subcortical brain. For the atlas database, the center of the cropping window was aligned with the center of mass of the subject's 7TAMIBrain segmentation masks, resulting in a set of N=60 images of equal size.

Here, a size of $192^3$ voxels was chosen to accommodate all brain sizes. Cropping of input images of patients and controls was performed similarly but after a fast direct segmentation by a non-linear registration process using the 7TAMIBrain template atlas.

3. This atlas database of N=60 subjects served as moving images during the MALF procedure. The entire procedure can be broken down into three main parts: (a) atlases to subject registrations, (b) atlases warping and label propagation, and (c) label fusion for subject segmentation.
   a. *Atlases-to-subject registrations*
      Registration between each atlas from 7TAMIBrain dataset and the target image (i.e., individual subject) was done using Advanced Normalization Tools' (ANTs) quick implementation of the symmetric image normalization method ('SyN') algorithm. Compared to the original implementation, the quick algorithm achieves a sustainable computing time by skipping the single-voxel scale. Comparisons between both methods, using Dice and volume similarity metrics, showed similar performances with a mean Dice score of 0.98 when comparing final segmentations.
   b. *Atlas warping and label propagation*
      The resulting set of transformations (i.e. affine matrix and a non-linear deformation field) were then applied to warp the atlases into each subject's space in the study cohort. As such, this step generated 60 warped atlases for every subject.
   c. *Label fusion*
      In this final step, label fusion was performed to construct – from the 60 atlases generated in b – a consensus atlas for each subject. A widely used and effective label fusion method is Majority Voting (MV), which finds the most common label across the 60 atlases for each voxel. However, since the computing of each label in the consensus atlas depends only on the average of the results obtained via the 60 registrations process, MV can be sensitive to registration biases. Therefore, the Joint Fusion (JF) algorithm implemented in ANTs' '*antsJointFusion*' was used instead which aims to reduce such biases. To increase computing speed, MV is still performed but only when agreement across 80% of references is reached. However, current parameters were set so that a full consensus was necessary to allow the use of MV instead of JF. This slows down the process, but the label fusion process has a small cost in computing time compared to step b. The resulting, final segmentations were used for further analyses.
