## Supplementary Results for "Multi-scale structural alterations of the thalamus and basal ganglia in focal epilepsy as demonstrated by 7T MRI"

#### Deformation and shape

##### Deformation

Compared to volume and $T_1$, tissue deformation shows inverse changes ($F_{2,76} = 6.567$, $p < .005$, Fig. S1A, top panel) with significant larger deformations for TLE patients compared to controls ($p < .01$). Larger deformations were consistently observed across nuclei ($F_{18,\,138} = 1.773$, $p < .05$; Wilk's $\Lambda = 0.660$, partial $\eta^2 = .19$, Fig. S1B, left heatmap) with strongest group effects seen for the central lateral and mediodorsal nuclei (both $p < .001$). In both cases both patient groups show larger averages compared to controls ($p < .05$). Ipsi- vs contralateral differences for whole thalamic and individual nuclei did not reach the significance threshold (Fig. S1B, right scatter plot and heatmap).

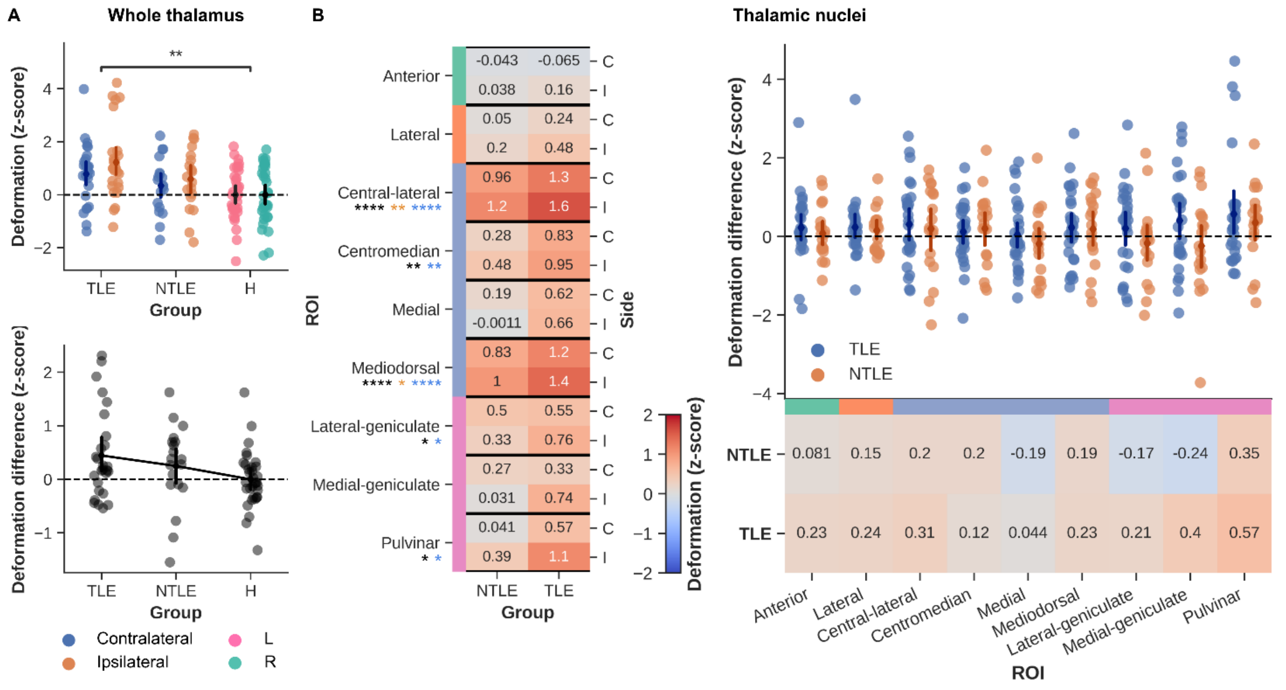


**Figure S1** – Deformation results. (A) Whole thalamic deformation (z-scored) was compared between groups (top panel) as well as between sides (i.e., ipsilateral-contralateral differences for patients, left-right for controls, bottom panel). (B) Similarly, average

deformation within each thalamic nuclei (z-scored) are shown for both patient groups (columns) and side (rows, left heatmap). Asterisks below nuclei labels indicate MANOVA (black), and pairwise comparison (orange for NTLE and blue for TLE, vs. controls) results. Strip plot and right heatmap in B illustrate the ipsilateral-contralateral deformation differences for each patient (color-coded by group). Color-bars alongside nuclei labels indicate group assignment. C = contralateral, I = ipsilateral, * p < .05, ** p < .01, *** p < .005, **** p < .001, Bonferroni corrected in case of multiple comparisons.

#### Shape

The surface-based shape metric constructed from vertex deformation, area and curvature properties, show increased averages for patients ($F_{2,76}$ = 19.46, p < .001) with strong increases for both patient groups (p < .001, Fig. S2, top panel). In particular, the shape differences for the ipsilateral side appeared higher for the TLE group (p < .05, Fig. S2, bottom panel). While ipsi- vs. contralateral differences remained undetected, thalamic nuclei were characterized by increased shape differences than controls ($F_{16, 138}$ = 2.622, p < .001; Wilk's $\Lambda$ = 0.588, partial $\eta^2$ = .23, Fig. S2B, left heatmap), in particular the anterior nucleus ($F_{2,76}$ = 8.758, p < .001).

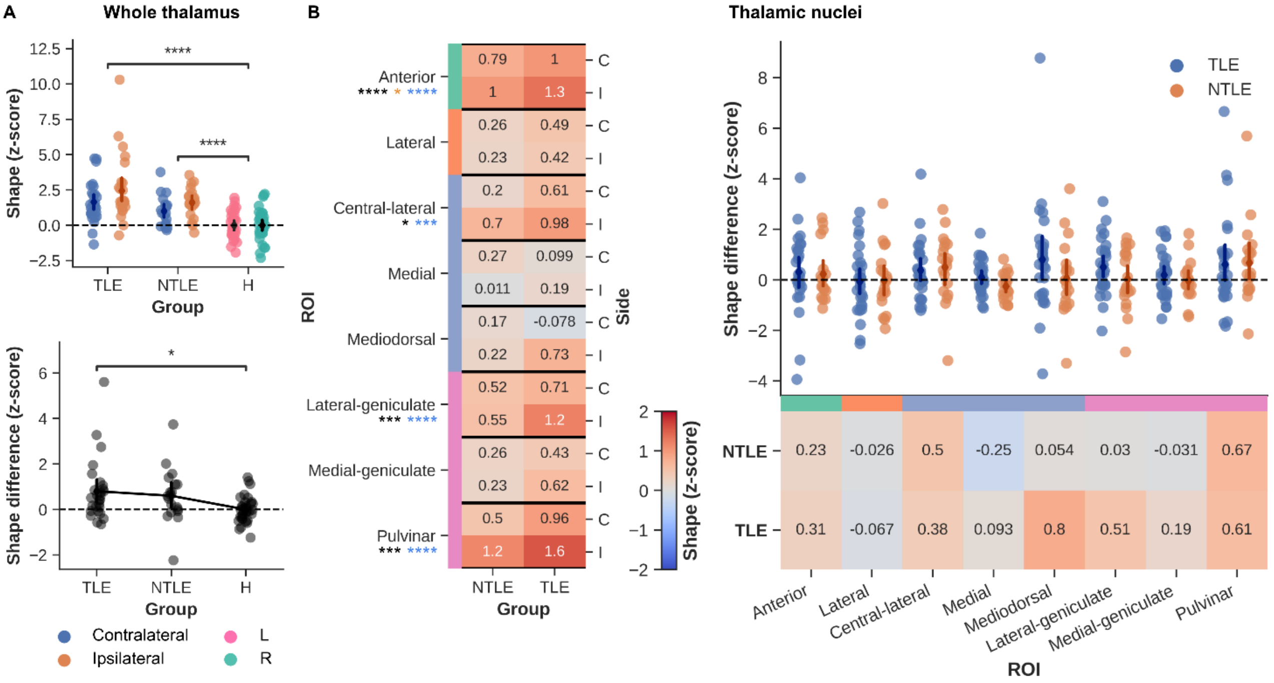


**Figure S2** – Shape results. (A) Whole thalamic shape (z-scored) was compared between groups (top panel) as well as between sides (i.e., ipsilateral-contralateral differences for patients, left-right for controls, bottom panel). (B) Similarly, average shape within each thalamic nuclei (z-scored) are shown for both patient groups (columns) and side (rows, left heatmap). Asterisks below nuclei labels indicate MANOVA (black), and pairwise comparison (orange for NTLE and blue for TLE, vs. controls) results. Strip plot and right heatmap in B illustrate the ipsilateral-contralateral shape differences for each patient (color-coded by group). Color-bars alongside nuclei labels indicate group assignment. C = contralateral, I = ipsilateral, * p < .05, ** p < .01, *** p < .005, **** p < .001, Bonferroni corrected in case of multiple comparisons.

### Replicating using THOMAS

To evaluate dependence of the findings on the thalamic atlas, main analyses were repeated using the THOMAS thalamic segmentation instead.

##### Comparison of 7T-AMI and THOMAS atlases

Fig. S3A shows a comparison between the 7T-AMI and THOMAS atlases when applied on the same subject. Differences exist in the number of segmented thalamic nuclei between the 7T-AMI and THOMAS atlases, most notably for the lateral and medial nuclei groups (in blue and pink, respectively, Fig. S3B). Whereas the lateral nucleus is treated as a single entity by the 7T-AMI atlas, the THOMAS atlas splits it up further into the ventral anterior (VA), ventral lateral posterior (VLp), ventral lateral anterior (VLa) and ventral posterior lateral (VPL) nuclei. In addition, the 7T-AMI atlas segments the medial and central lateral nuclei (along the internal medullary lamina) while these are not considered by THOMAS.

Dice Similarity Coefficients (DSCs) were then calculated to quantify the overlap between each nucleus from both atlases for a single subject. DSC values were than averaged across subjects and plotted in Fig. S3C. Here, the pulvinar (0.84) and mediodorsal (0.78) nuclei show highest overlap, while it appears more fractioned for the central-lateral and lateral nuclei.

In general, whole thalamic volume (z-scored) from both atlases (with both hemispheres pooled) showed a high correlation across controls, $r(64) = .84$, $p < .001$ (Fig. S3D, left panel). This was also true for the individual nuclei (i.e., those shared between atlases, $p_{FDR} < .001$, middle panel), except for the lateral geniculate nucleus (LGN, $p_{FDR} > .05$). Left-right volume differences follow the same trend with relative high correspondence between atlases, $r(64) = .61$, $p < .001$.

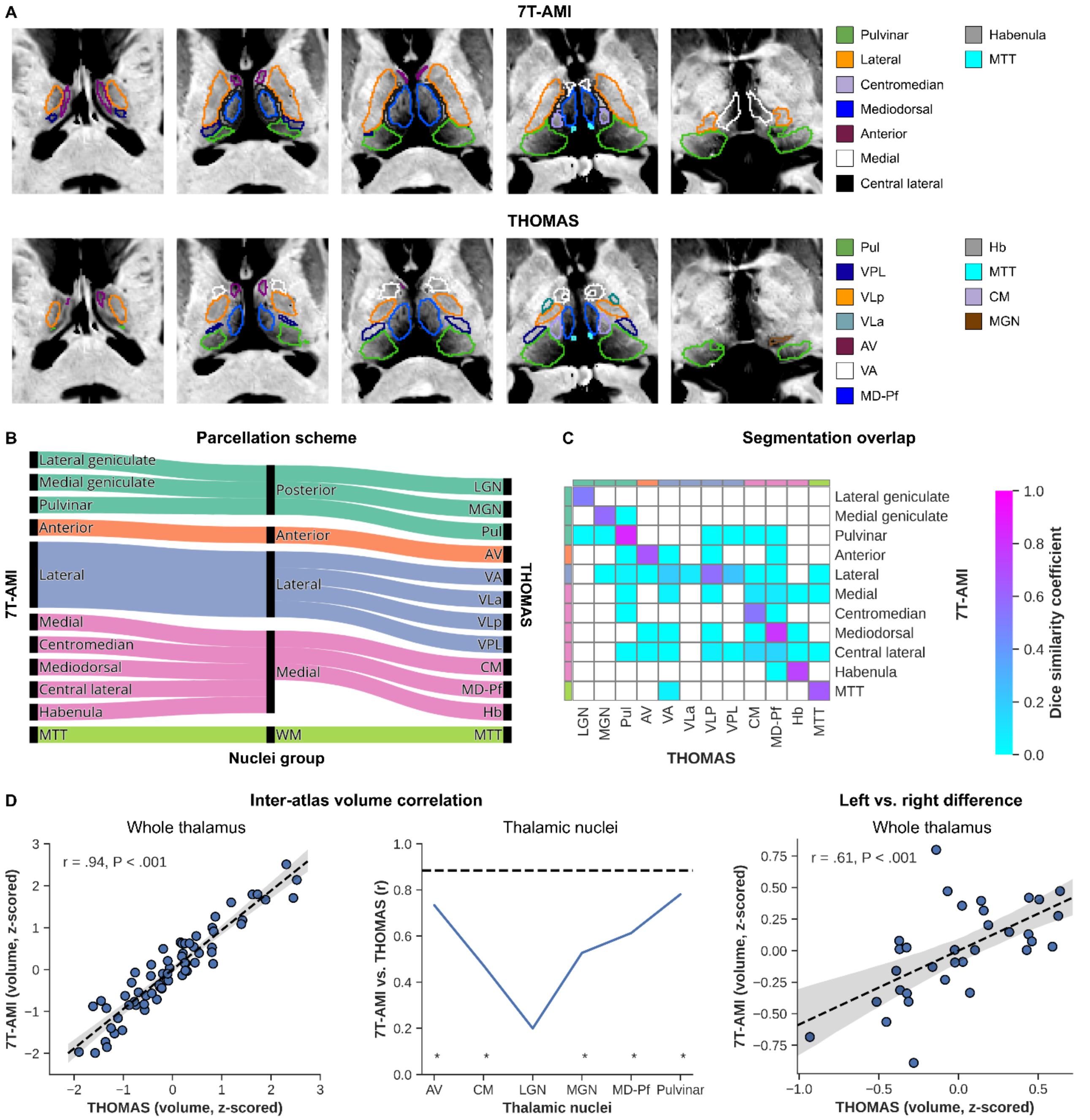


**Figure S3** – Comparison of 7T-AMI and THOMAS thalamic atlases. (A) Montages of axial slices show both atlases (rows) when applied to the same example subject. Note that not all nuclei are visible on these slices. (B) Sankey plot showing correspondence between both atlas nomenclatures. Center nodes indicate color-coded grouping of individual nuclei. (C) Spatial overlap between each atlas' nuclei based on the Dice Similarity Coefficient. (D) Correlation analyses between both atlases when extracting whole thalamic volume (left), nuclei volumes (center, asterisks indicating significant correlation) and left vs. right whole thalamus volume differences (right).

#### Volume

Whole thalamic volume measures using THOMAS follow the same trend as in Fig. 3 based on the 7T-AMI atlas ($F_{2,76} = 11.37$, $p < .001$) with significantly decreased volume for TLE and NTLE patients compared to controls (both $p < .001$, Fig. S4A). In contrast, ipsilateral volume appears significantly reduced in TLE ($p < .05$) as well as NTLE ($p < .01$) patients compared to controls using the THOMAS atlas.

Nuclei-wise, reduced volumes were observed for the patient groups ($F_{20, 134} = 2.454$, $p < .005$; Wilk's $\Lambda = 0.536$, partial $\eta^2 = .27$, Fig. S4B). Similarly, the strongest effect was observed for the MD-Pf nucleus ($F_{2,76} = 12.609$, $p < .001$), with both TLE and NTLE showing reduced volumes compared to controls ($p < .001$), and weakest effect for the AV nucleus (N.S.). There was no systematic ipsi- vs. contralateral volume difference between groups ($F_{20, 134} = 1.538$, $p = .078$; Wilk's $\Lambda = 0.661$, partial $\eta^2 = .19$).

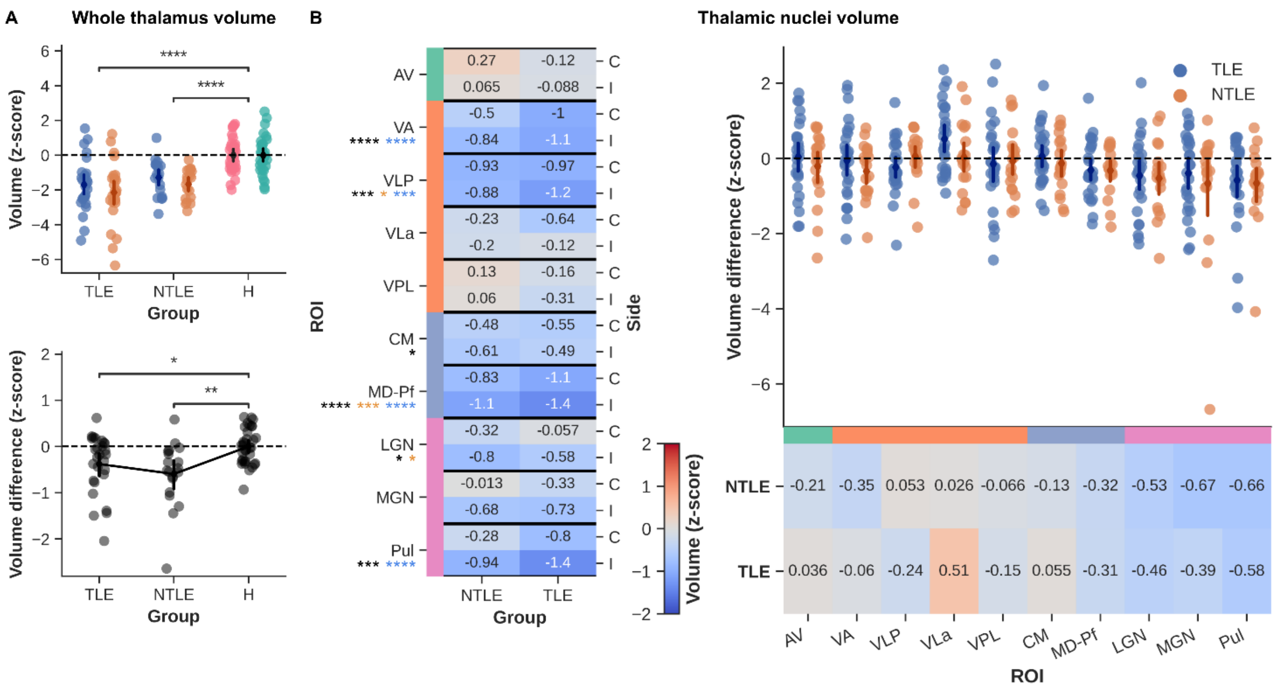


**Figure S4** – Volumetric results. (A) Whole thalamic volume (z-scored) was compared between groups (top panels) as well as between sides (i.e., ipsilateral-contralateral differences for patients, left-right for controls, bottom panels). (B) Similarly, average volumes for thalamic nuclei (z-scored) are shown for both patient groups (columns) and side (rows, left heatmaps). Asterisks below nuclei labels indicate MANOVA (black), and pairwise comparison (orange for NTLE and blue for TLE, vs. controls) results. Strip plots and right heatmaps in B illustrate the ipsilateral-contralateral $T_1$ differences for each patient (color-coded by group). Color-bars alongside nuclei labels indicate group assignment. C = contralateral, I = ipsilateral, * $p < .05$, ** $p < .01$, *** $p < .005$, **** $p < .001$, Bonferroni corrected in case of multiple comparisons.

## $T_1$

As for volume, thalamic $T_1$ was significantly different across groups ($F_{2,76} = 4.455$, $p < .05$), and in particular decreased for the TLE ($p < .05$, Fig. S5A) and not NTLE patients. This general group effect extends towards the individual nuclei ($F_{20, 134} = 2.597$, $p < .005$; Wilk's $\Lambda = 0.519$, partial $\eta^2 = .28$, Fig. S5B). Here, the VPL nucleus was most strongly affected with reduced $T_1$ values observed for both groups ($p < .001$) while the AV nucleus appeared least impacted (N.S.). In line with the other findings, systematic ipsi- vs. contralateral $T_1$ differences between groups remained absent and were therefore not explored at the individual nuclei level.

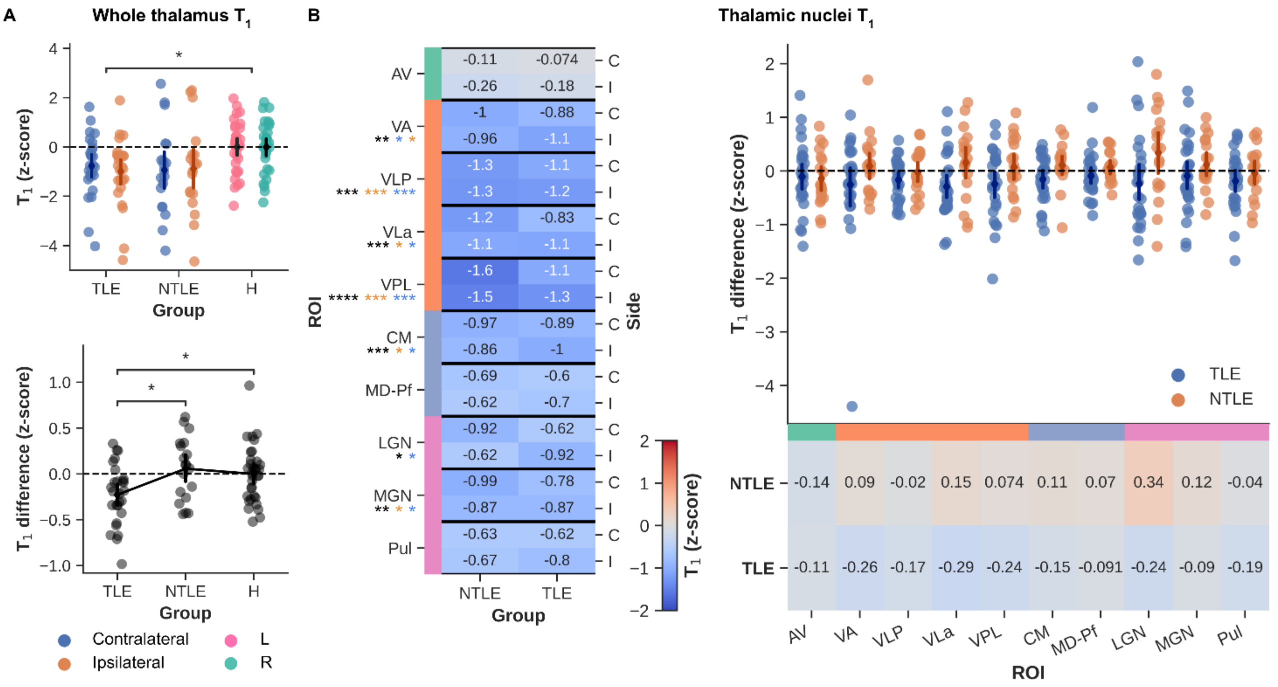


**Figure S5** – $T_1$ results. (A) Whole thalamic $T_1$ (z-scored) was compared between groups (top panels) as well as between sides (i.e., ipsilateral-contralateral differences for patients, left-right for controls, bottom panels). (B) Similarly, average $T_1$ for thalamic nuclei (z-scored) are shown for both patient groups (columns) and side (rows, left heatmaps). Asterisks below nuclei labels indicate MANOVA (black), and pairwise comparison (orange for NTLE and blue for TLE, vs. controls) results. Strip plots and right heatmaps in B illustrate the ipsilateral-contralateral $T_1$ differences for each patient (color-coded by group). Color-bars alongside nuclei labels indicate group assignment. C = contralateral, I = ipsilateral, * $p < .05$, ** $p < .01$, *** $p < .005$, **** $p < .001$, Bonferroni corrected in case of multiple comparisons.

#### Deformation

The extent of tissue deformation differed significantly across groups ($F_{2,76} = 5.674$, $p < .01$) with larger values observed for TLE patients ($p < .01$), and in particular for the ipsilateral side in these patients ($p < .05$) compared to controls. A similar group effect was seen across individual nuclei ($F_{20,134} = 1.608$, $p = .05$; Wilk's $\Lambda = 0.650$, partial $\eta^2 = .19$, Fig. S6B), without modulation by side. Deformation was especially high in the MD-Pf ($F_{2,76} = 10.38$, $p < .001$) and CM ($F_{2,76} = 6.217$, $p < .005$) nuclei.

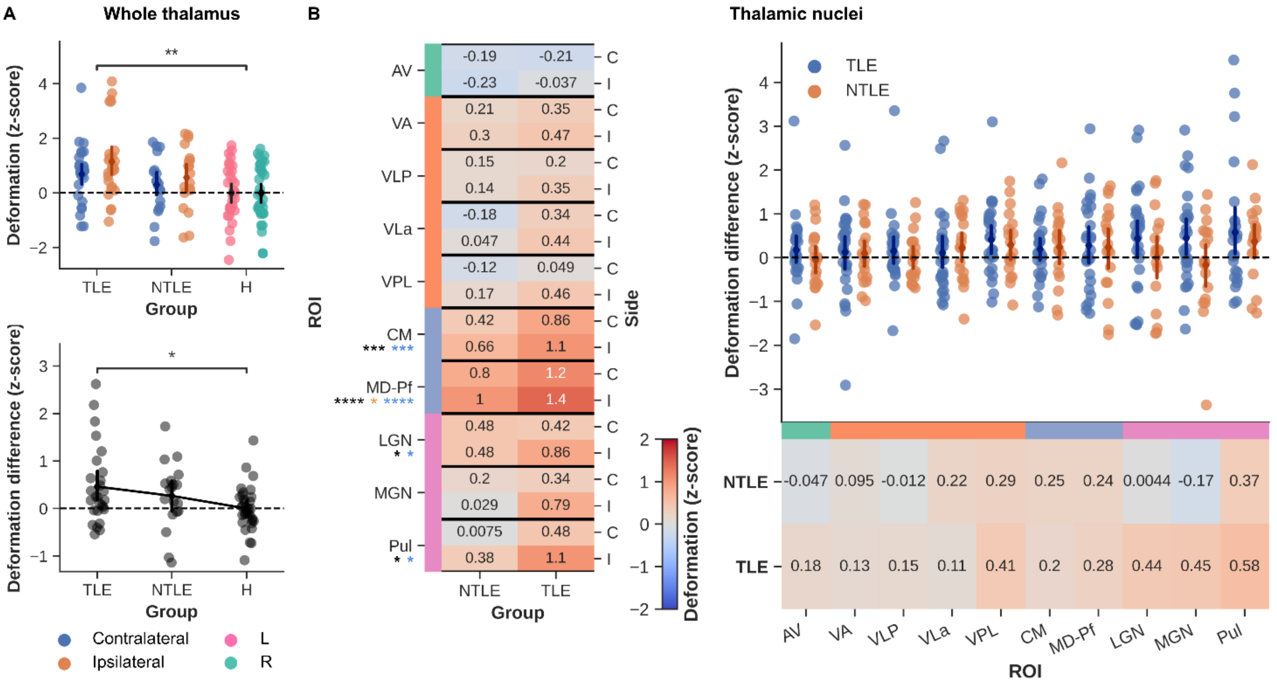


**Figure S6** – Deformation results. (A) Whole thalamic deformation (z-scored) was compared between groups (top panel) as well as between sides (i.e., ipsilateral-contralateral differences for patients, left-right for controls, bottom panel). (B) Similarly, average deformation within each thalamic nuclei (z-scored) are shown for both patient groups (columns) and side (rows, left heatmap). Asterisks below nuclei labels indicate MANOVA (black), and pairwise comparison (orange for NTLE and blue for TLE, vs. controls) results. Strip plot and right heatmap in B illustrate the ipsilateral-contralateral deformation differences for each patient (color-coded by group). Color-bars alongside nuclei labels indicate group assignment. C = contralateral, I = ipsilateral, * p < .05, ** p < .01, *** p < .005, **** p < .001, Bonferroni corrected in case of multiple comparisons.

#### Shape

Whole thalamic shape follows the same, but inverse relationship with group as observed for volume ($F_{2,76}$ = 10.381, p < .001). Averaged across the thalamus, both patient groups show increased shape values compared to controls (both p < .01). This effect holds across the individual nuclei across the nuclei ($F_{20,134}$ = 1.785, p < .05; Wilk's Λ = 0.623, partial η2 = .21, Fig. S7B) but the distribution of these changes differs with respect to volume. Here, strongest group effects were observed for the AV ($F_{2,76}$ = 7.876, p < .001) and pulvinar ($F_{2,76}$ = 5.433, p < .01) regions. No differences were observed between ipsi- and contralateral sides here as well.

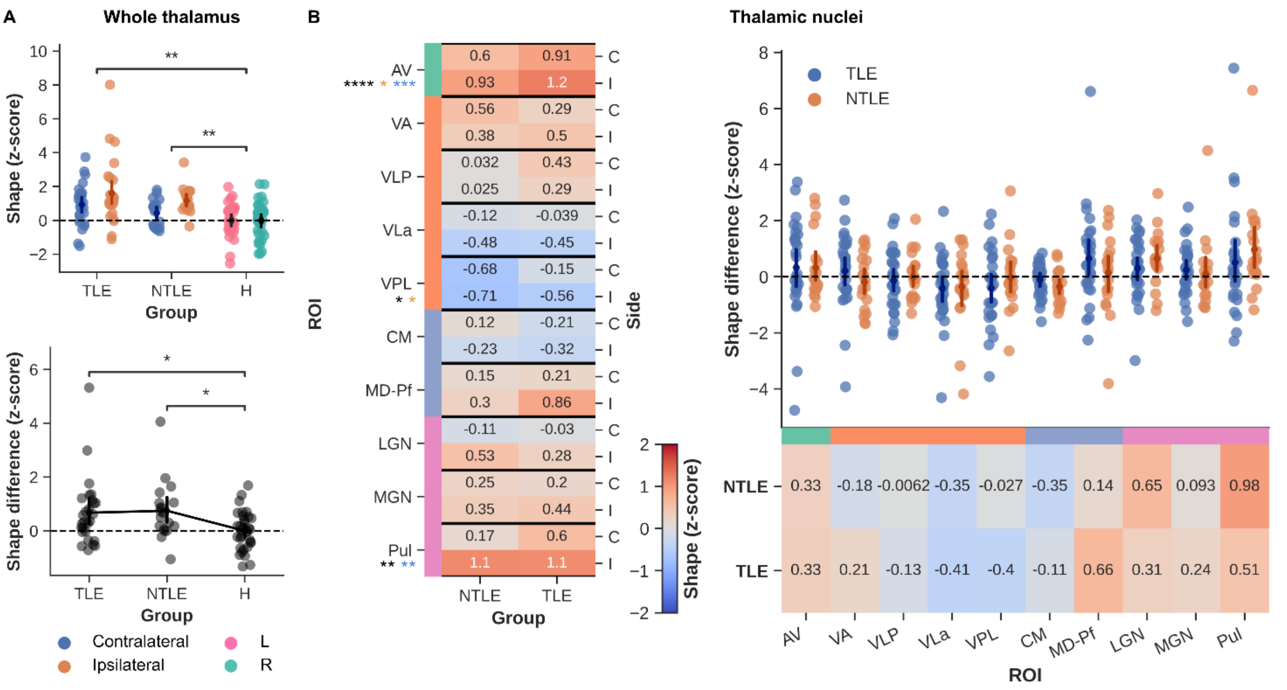


**Figure S7** – Shape results. (A) Whole thalamic shape (z-scored) was compared between groups (top panel) as well as between sides (i.e., ipsilateral-contralateral differences for patients, left-right for controls, bottom panel). (B) Similarly, average shape within each thalamic nuclei (z-scored) are shown for both patient groups (columns) and side (rows, left heatmap). Asterisks below nuclei labels indicate MANOVA (black), and pairwise comparison (orange for NTLE and blue for TLE, vs. controls) results. Strip plot and right heatmap in B illustrate the ipsilateral-contralateral shape differences for each patient (color-coded by group). Color-bars alongside nuclei labels indicate group assignment. C = contralateral, I = ipsilateral, * $p < .05$, ** $p < .01$, *** $p < .005$, **** $p < .001$, Bonferroni corrected in case of multiple comparisons.

#### Partial Least Squared (PLS) analyses

In line with the mean-centered PLS findings utilizing the 7T-AMI thalamic segmentation, the THOMAS atlas reveals similar LVs when contrasting NTLE, TLE and controls data or epilepsy types (Fig. S8). For the former, the LV (explaining 75.9% of the variance, $p_{perm} < .001$, Fig. S8A) shows a similar impact of metric type ($F_{3,112} = 10.25$, $p < .001$) with predominant contributions from volume information. In addition, structure (i.e., thalamus vs other basal ganglia, $F_{1,112} = 10.39$, $p < .005$) as well as side (ipsi- vs. contralateral, $F_{1,112} = 9.977$, $p < .005$) have a comparable effect. Here, non-thalamic as well as ipsilateral features (both $p < .05$) appeared more impactful.

When contrasting across epilepsy types (63.6% explained variance, $p_{perm} < .05$, Fig. S8B) distribution of feature contributions shifted towards $T_1$-based measures ($p < .001$ compared to shape and volume) while reliability of thalamic features increases with respect to the other basal ganglia ($p < .01$). This is in line with the 7T-AMI findings.

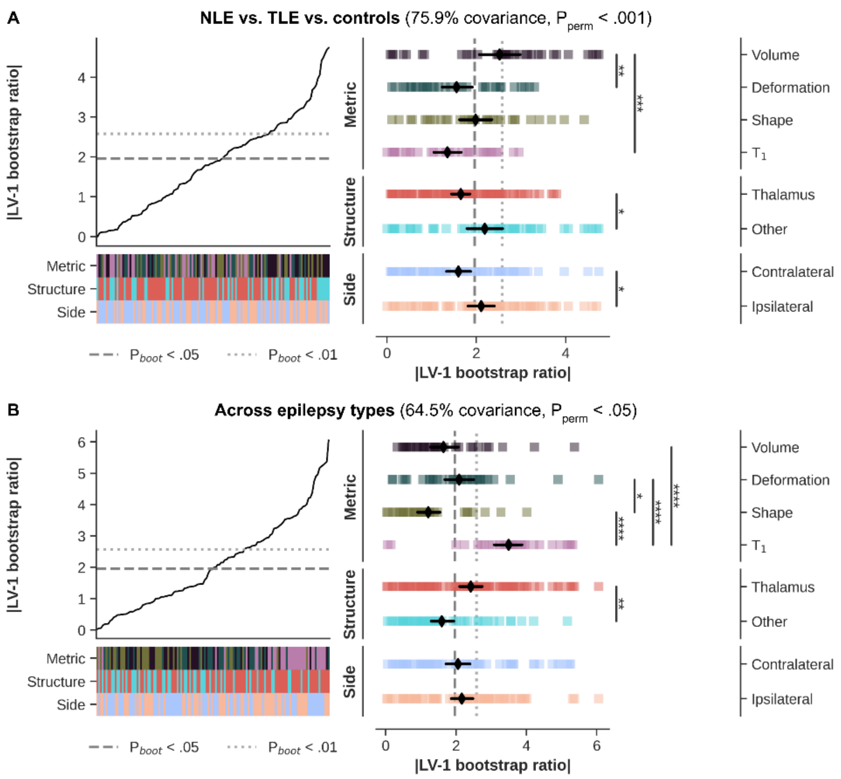


**Figure S8** – Mean-centered PLS results. (A) Line plot shows sorted bootstrapped ratios when contrasting NTLE, TLE and controls. Heatmaps indicate feature properties (metric type, structure, and side, color-coded as in Fig. 2). Strip plot shows the decomposition of features based on feature properties with black diamonds indicating average BSR (± 95% CI). (B) Same as A but contrasting epilepsy types. * p < .05, ** p < .01, *** p < .005, **** p < .001, Bonferroni corrected in case of multiple comparisons.

No major differences with 7T-AMI were observed with regards to the behavioral PLS results (Fig. S9A). The latent variable (35.2% explained variance, $p_{perm} < .05$) encompassed a comparable ranking of MRI-based features. Here, BSRs varied as function of metric type ($F_{3,112} = 16.48$, $p < .001$) and side ($F_{1,112} = 12.73$, $p < .005$). For the clinical features, both nr. of epileptic zones and lesion detected show strongest correlation with the latent variable (both $p_{boot} < .05$). The scatter plots in Fig. S9C show exemplifies the relationship between MRI features (z-scored) and the two clinical features showing the strongest correlation in B.

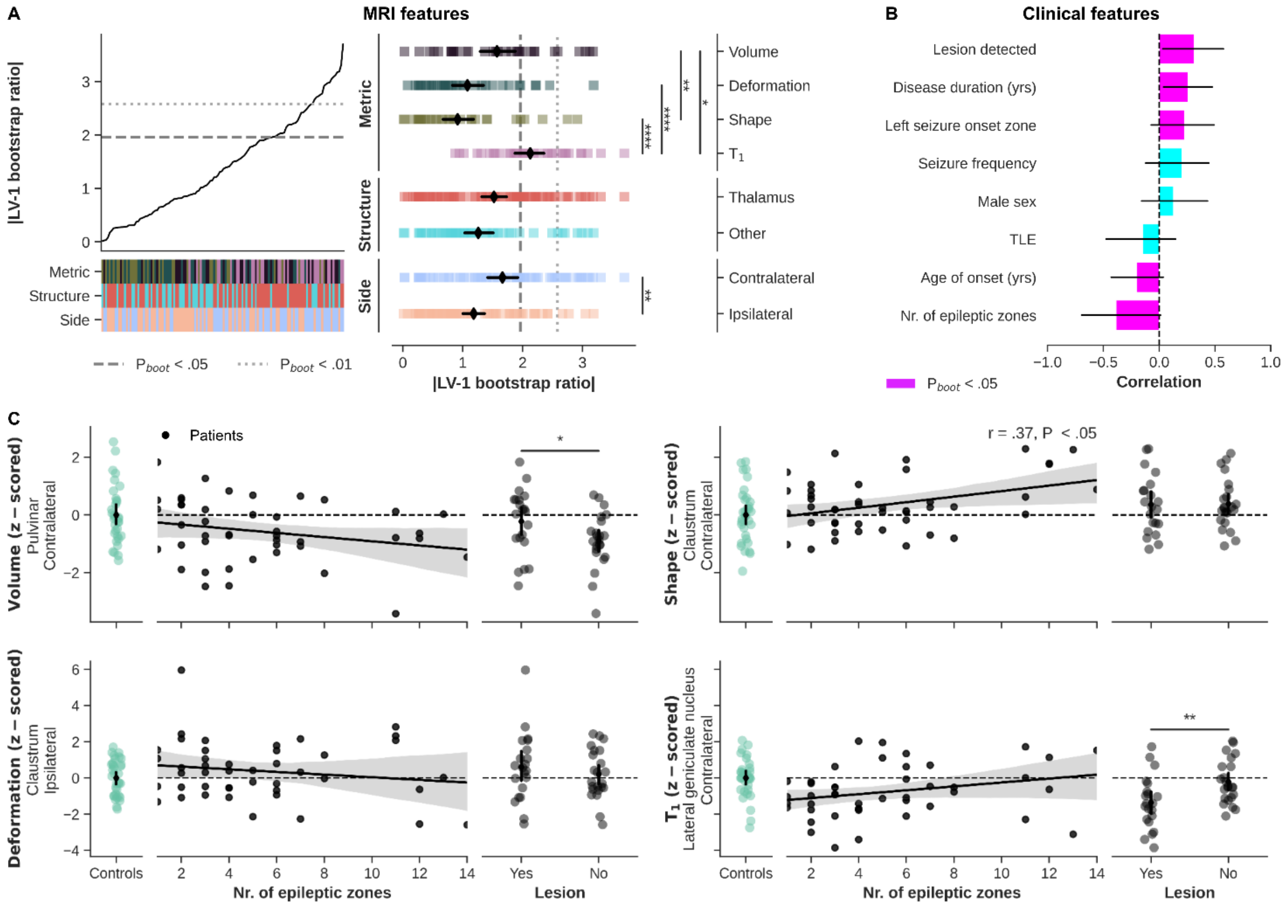


**Figure S9** – Behavioral PLS results. (A) Line plot shows sorted bootstrapped ratios with heatmaps indicating feature properties (metric type, structure, and side, color-coded as in Fig. 2). Strip plot shows the decomposition of features based on feature properties with black diamonds indicating average BSR (± 95% CI). (B) Ranking of clinical features in terms of correlation with the LV. (C) Correlation between clinical features and MRI-based z-scores. * $p < .05$, ** $p < .01$, *** $p < .005$, **** $p < .001$, Bonferroni corrected in case of multiple comparisons.
